# A comprehensive atlas of testicular interstitium reveals Cd34^+^/Sox4^+^ mesenchymal cells as potential Leydig cell progenitors

**DOI:** 10.1101/2024.08.02.606288

**Authors:** Xiaojia Huang, Kai Xia, Meiling Yang, Mengzhi Hong, Meihua Jiang, Weiqiang Li, Zhenmin Lei, Andy Peng Xiang, Wei Zhao

## Abstract

The declining rates of male fertility pose a significant clinical challenge, primarily due to our limited understanding of the testicular interstitium, which is crucial for male reproductive health. Here, we conducted a comprehensive analysis of the single-cell transcriptomic landscape of the murine testicular interstitium across the postnatal lifespan. Our investigation unveiled a previously unrecognized population of Cd34^+^/Sox4^+^ mesenchymal cells nestled within the interstitium, hinting at their potential as Leydig cell progenitors. During the aging process of Cd34^+^/Sox4^+^ mesenchymal cells, we observed a decline in glutathione levels within the testicular interstitium. Remarkably, these Cd34^+^/Sox4^+^ mesenchymal cells exhibited clonogenic self-renewal capacity and an impressive propensity to differentiate into Leydig cells. Intriguingly, when transplanted into Leydig cell-disrupted or failure models, Cd34^+^/Sox4^+^ cells efficiently colonized the testicular interstitium, resulting in a notable increase in testosterone production. Exploring the epigenetic landscape, we identified critical transcription factors, most notably Sox4, governing the stem cell fate of Cd34^+^/Sox4^+^ mesenchymal cells. Overall, this comprehensive reference atlas of lifespan testicular Leydig cells presents significant findings that may guide the development of cell-based strategies for treating testicular hypogonadism in elderly individuals.

## Introduction

Decreasing fertility rates has emerged as a significant social and medical concern, with male infertility accounting for at least half of the cases. Conventional semen analysis offers restricted prognostic utility for male fertility, largely because a considerable number of male infertility incidents are attributed to dysfunctions in the testicular interstitium. Among the testis interstitial cells, Leydig cell (LC) (Haider, 2004) , play a pivotal role in regulating male fertility(Ge et al., 2008; Potter and DeFalco, 2017; Zirkin and Papadopoulos, 2018). Throughout testicular development, LC are responsible for testosterone production, a hormone crucial for the fetal differentiation of the male reproductive tract, the virilization of male external genitalia, and the pubertal maturation of Sertoli cells. In adult males, LC continue to express androgens, which are essential for maintaining masculine characteristics, including sex drive, erectile function, and fertility(Forest, 2008). Despite the critical importance of LC to male fertility, there is a notable gap in our understanding of their temporal dynamics and heterogeneity during the age-related decline in reproductive capability.

Prior research has delineated two distinct populations of LC populations within the testicular interstitium: fetal LC (FLC) and adult LC (ALC)(DeFalco et al., 2011). FLC, emerging within the fetal testes interstitium, reach their numerical apex around the time of birth. While earlier studies posited the absence of FLC in postnatal mouse testes, advancements in lineage-tracing technologies have since unveiled that FLC undergo significant regression post-birth, with only a minimal subset persisting into adulthood(Liu et al., 2016; Morohashi et al., 2015; Shima et al., 2018; Wen et al., 2016). In contrast, ALC manifest at puberty, occupying the testicular interstitium to sustain male secondary sexual characteristics and fertility through adult life. Characterized as terminally differentiated, ALC possess a restricted capacity for regeneration and proliferation, rendering them susceptible to age-related dysfunctions(Chamindrani Mendis-Handagama and Siril Ariyaratne, 2001; Midzak et al., 2009). Consequently, elucidating the molecular evolution of LC across the lifespan is imperative for devising effective interventions aimed at preserving male fertility and addressing age-associated testicular dysfunctions.

To acquire an in-depth comprehension of the differentiation and ongoing maintenance of ALC across postnatal development, concerted research efforts have been focused on pinpointing the stem/progenitor cell reservoirs from which ALC derive. Prior investigations, inclusive of our own, have established Nestin as a crucial biomarker for stem LC situated in the peritubular regions of seminiferous tubules and within perivascular niches(Jiang et al., 2014; Zang et al., 2017). Notably, recent studies underscore the critical importance of a finely tuned equilibrium between the generation of reactive oxygen species (ROS) and the capacity of intracellular antioxidant defenses in sustaining optimal testosterone synthesis(Ding et al., 2016). Perturbations in this equilibrium can lead to significant physiological repercussions for LC, notably a reduction in testosterone output. Thus, it is critical to persist in exploring the variety of LC, seeking to elucidate the complex molecular interactions between LC and their surrounding microenvironment.

Recent advancements in single-cell RNA sequencing (scRNA-seq) have illuminated the molecular alterations accompanying aging in testicular tissue across both human and murine models. For instance, research by Nie et al. uncovered age-associated regulatory changes impacting spermatogenesis(Chen et al., 2021). Another investigation revealed an age-induced imbalance between undifferentiated and differentiated spermatogonia stem cells(Hermann et al., 2018). Although these studies have enhanced our understanding of the aging process within the testes, there remains a vital need for further exploration into the molecular changes occurring at various postnatal stages (neonatal, pre-pubertal, adolescent, adult, and aged phases) within the testicular interstitium.

In this study, we utilized scRNA-seq to delve into the molecular diversity and temporal patterns of LC within the testicular interstitium across postnatal development. Our analyses revealed a hitherto unidentified population of Cd34^+^/Sox4^+^ mesenchymal cells, pinpointed as key progenitors instrumental in the maintenance and renewal of LC. Additionally, we noted an age-associated functional decline in glutathione metabolism. Subsequent in vitro and in vivo experiments underscored the regenerative capabilities of these Cd34^+^/Sox4^+^ mesenchymal cells within the testicular interstitium. We further identified the critical role of the transcription factor Sox4 in maintaining the stemness of Cd34^+^/Sox4^+^ mesenchymal cells. Our results pave the way for the development of improved therapeutic strategies for male infertility and the enhancement of reproductive health outcomes.

## Results

### Single-cell transcriptomic analysis of testicular interstitium unveils LC heterogeneity

To enrich for interstitial cells, we employed flow cytometry on our samples, maximizing the collection of interstitial cells (Figure 1—figure supplement 1a). We harvested cells from mouse testes at key developmental milestones: neonatal (1 week), adolescence (1 month), adult (2 months), middle-aged (8 months), and aged (24 months). Utilizing the 10x Genomics Chromium platform, we conducted single-cell RNA sequencing (scRNA-seq) on samples from each of these stages (Figure 1a). Through uniform manifold approximation and projection analysis, we delineated five distinct cell types characterized by signature genes identified in prior research: spermatogenic cell (Prm1, Oaz3, Dbil5), Sertoli cell (Wt1, Sox9, Amh), immune cell (Cd45, Cd68, Cd74), mesenchymal cell (Dcn, Col1a2, Col3a1), and LC (Nr5a1, Cyp17a1, Hsd3b1) (Figure 1—figure supplement 1b and c).

**Figure 1.**
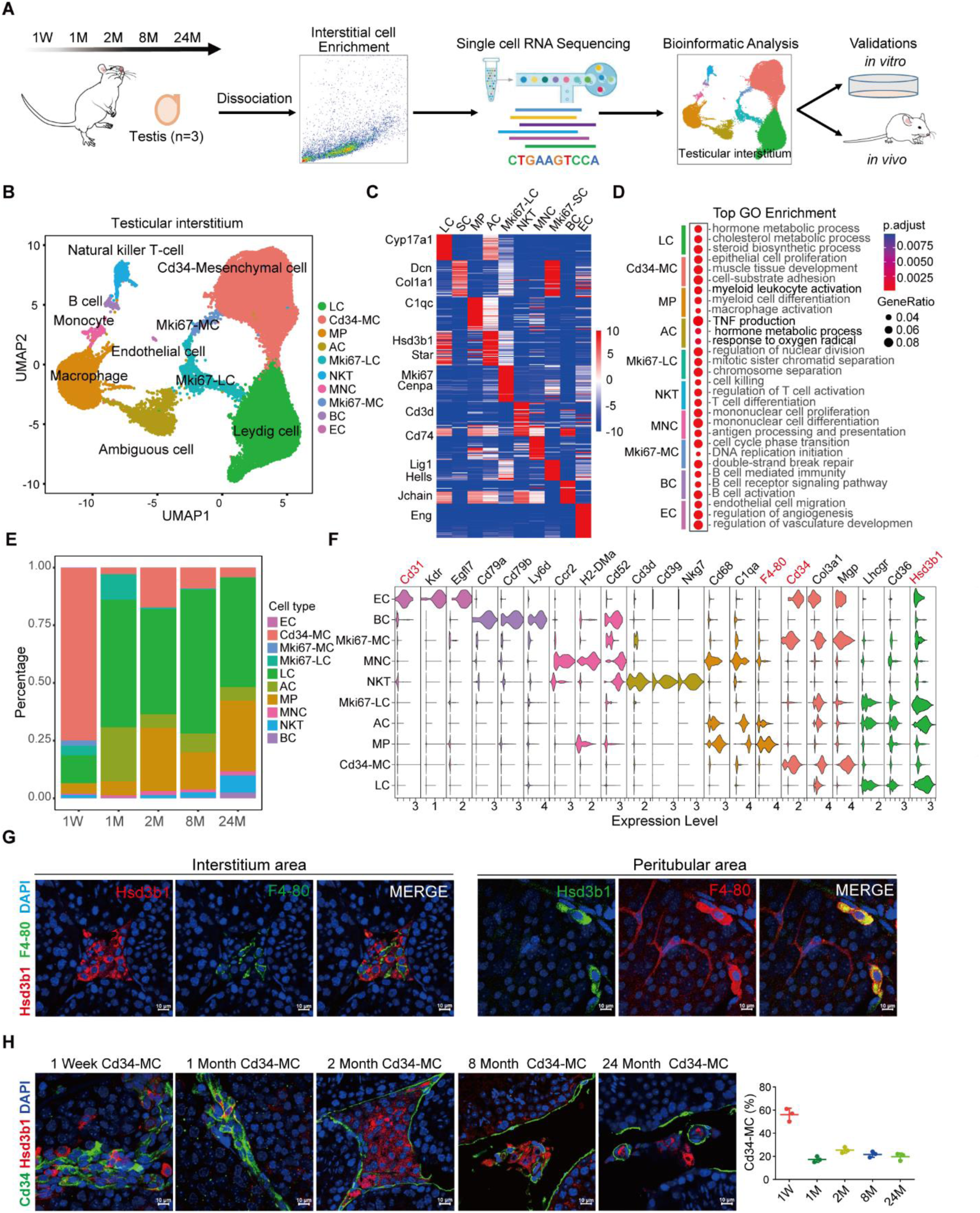
Characterization of cell populations in the mouse testicular interstitium. (**a**) Overview of the experimental design and workflow employed for single-cell RNA sequencing (scRNA-seq) to explore cell populations in the testicular interstitium at various developmental stages: neonatal, adolescent, adult, middle-aged and aged, with three replicates per stage. (**b**) Uniform manifold approximation and projection (UMAP) visualization illustrating the diversity of cell types identified within the testicular interstitium across the developmental spectrum: neonatal to aged stages in mice. (**c**-**d**) Heatmap (**c**) showcasing the top ten differentially expressed genes and dot plot (**d**) highlighting the predominant GO terms associated with each cell cluster. (**e**) Bar graphs showing the distribution of single-cell proportions at different developmental stages within the sample set. (**f**) Violin plots elucidating expression trends of specific markers crucial for delineating primary cell types in the testicular interstitium. (**g**) Immunofluorescence images of testicular tissue double-labeled for Hsd3b1 and F4-80, with cells co-expressing these markers (Ambiguous cells, AC) indicated in yellow. Scale bar: 10 µm. (**h**) Immunofluorescence staining for Hsd3b1 and Cd34 in testicular tissue to demarcate Leydig cells (LC) and mesenchymal cells (MC), respectively. Scale bar: 10 µm.

Our comprehensive focus across the interstitium uncovered ten distinct populations of interstitial cells, including well-characterized types such as LC, mesenchymal cell (MC), endothelial cell (EC), and various immune cells-macrophage (MP), monocyte (MNC), Natural killer T lymphocyte (NKTC), and B lymphocyte (BC) (Figure 1b, Figure 1—figure supplement 1d and e). Additionally, we identified two proliferating cell populations: Mki67-LCs, marked by elevated expression of mitosis-related genes (Mki67, Cenpa) alongside LC signatures (Hsd3b1, Star); and Mki67-MCs, characterized by DNA replication-related gene expression (Hells, Lig1) and mesenchymal cell signatures (Dcn, Col1a2) (Figure 1b-d).

Notably, our analysis identified a distinct cell type displaying gene expression profiles characteristic of both macrophages and LC, which we termed Ambiguous cell (AC) (Figure 1c). Gene Ontology (GO) analysis of differentially expressed genes (DEGs) linked AC with processes such as TNF production, hormone metabolism, and response to oxygen radicals (Figure 1d). Intriguingly, AC populations proliferate rapidly during adolescence before declining to stabilize at a lower proportion in later life stages (Figure 1e). Comparative analysis between AC and LC expression profiles showed AC-specific upregulation of DEGs involved in the positive regulation of responses to external stimuli (Figure 1—figure supplement 1f and g). Moreover, when comparing AC expression patterns to those of macrophages, we found an enrichment of AC-specific upregulated DEGs in hormone metabolic pathways (Figure 1—figure supplement 1h and i).

Aging correlated with significant increases in immune cell populations, such as macrophage, B lymphocyte, and T lymphocyte (Figure 1e). Critically, we identified a previously unknown population, Cd34-MC, which was abundant during neonatal stages but gradually declined with development (Figure 1e). Utilizing their specific surface markers (Figure 1f), immunofluorescence staining was employed to confirm the presence and location of AC (Figure 1g) as well as a marked reduction in Cd34-MC post-puberty (Figure 1h).

We next explored the temporal transcriptional shifts within the immune-related cells of the testicular interstitium, starting from the postnatal stage (Figure 1—figure supplement 2a). Macrophages, which are notably prevalent in the adult testis interstitium, displayed significant changes in their gene expression profiles over time (figure supplement 2b, c and d). CellChat analyses indicated consistent engagement between macrophages with LCs and Cd34-MC1 cells via the VCAM pathway across all five studied stages hinting at VCAM1’s potential role in enhancing adhesion or facilitating macrophage migration to the testis. Notably, the activity of the VCAM pathway decreased significantly by 24 months. Additionally, macrophages were found to secrete IGF1 and Visfatin (Figure 1—figure supplement 2e), which targeted LCs and likely promoted testicular steroidogenesis (Hameed et al., 2012; Jeremy et al., 2017; Lin et al., 1986; Ocón-Grove et al., 2010).

### Dynamic gene expression patterns in interstitial cells from infancy to adulthood

Clustering analysis exposed a continuous grouping among LC, AC, and MC, leading to a focused re-clustering of LC and MC across all ages. This refined analysis delineated seven distinct sub-clusters, corroborating previous research and recognized markers: Cd34-MC, Mki67-MC, primary LC (PLC, marked by Hsd3b6, Cyp51), immature LC (ILC, marked by Cyp17a1), mature LC (MLC, marked by Hsd3b1), Mki67-positive LC (Mki67-LC), Mki67-positive mesenchymal cell (Mki67-MC), and ambiguous cell (AC) (Figure 2a, Figure 2—figure supplement 3a). Furthermore, cell-type-specific markers were also unveiled, including Lcn2 and Ptgds for ILC, Cyp51 and Ass1 for PLC, and Kcnk3 and Agt for MLC (Figure 2b). A heatmap showcasing the top 10 DEGs and three representative markers are shown (Figure 2—figure supplement 3b). GO analysis of these DEGs provides insights into their roles in cellular functions (Figure 2—figure supplement 3c). Investigation of the differentiation trajectories for LC and MC via expression patterns of testosterone-synthesis genes indicated a progressive enhancement consistent with the proposed differentiation pathway (Figure 2—figure supplement 3d and e), suggesting that LC may derive from MC while exhibiting distinct functional characteristics.

**Figure 2.**
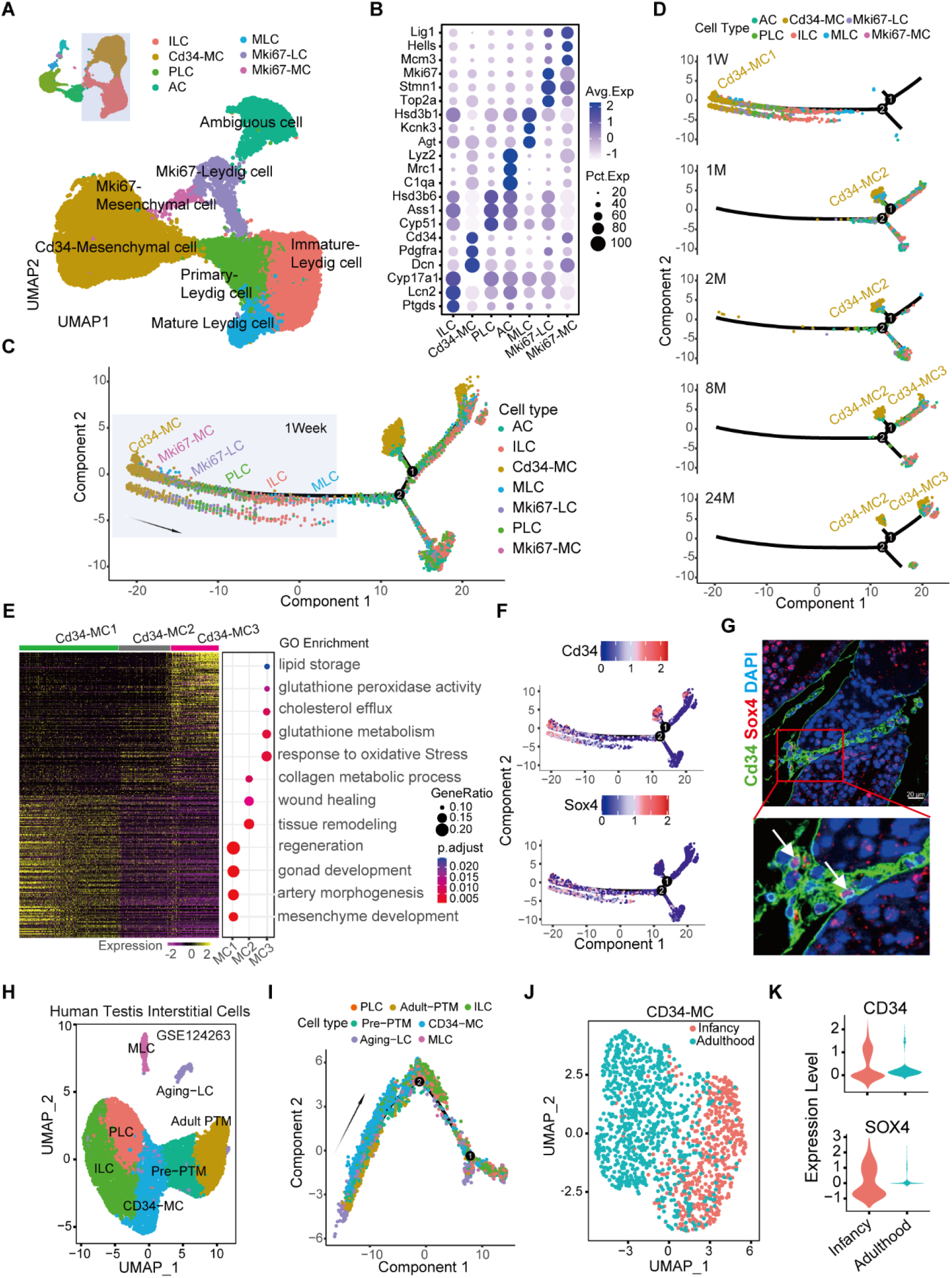
Dynamic gene-expression patterns of LCs subtypes at stepwise developmental stages. (**a**) UMAP analysis delineates LC and MC subtypes across developmental stages: neonatal (1 week), adolescent (1 month), adult (2 months), middle-aged (8 months), and aged (24 months) in mouse testes. (**b**) Dot plot showcases differentially expressed genes (DEGs) across LC subtypes. (**c**) Pseudotime trajectory, constructed with Monocle, traces the developmental lineage of LC subtypes. (**d**) Deconvolution plot demonstrating the gradual change of subtypes from neonatal to aged stages and from Progenitor (Cd34-MC) to Mature Leydig cell (MLC) as predicted by the pseudotime trajectory. (**e**) Heatmap and corresponding dot plot reveal DEG patterns within three Cd34-MC subgroups and their top GO enrichments, respectively. (**f**) Pseudotime trajectory plot illustrating the expression profiles of genes that become restricted to Cd34-MC1. (**g**) Immunofluorescence staining of Cd34 and Sox4 within the testicular interstitium at 3 weeks. Scale bar: 20 µm. (**h**) UMAP visualization of cell types within the testicular interstitium at infant and adult human stages (data sourced from GSE124263). (**i**) Pseudotime traces the developmental path of interstitial cells, color-coding each cell by subtype. (**j**) UMAP plot illustrating the presence of human CD34-MC subtype cells within the testicular interstitium during both infant and adult stages of development. (**k**) Violin plots depict CD34 and SOX4 expression in human CD34-MC cells at both infant and adult developmental stages.

To further investigate the developmental relationship and differentiation paths within LC lineages, we performed a comprehensive analysis spanning their developmental trajectory over time. Pseudotime trajectory analysis revealed marked variations in gene expression profiles between early postnatal (1-week-old) samples and those from adult stages, with the Cd34-MC population diverging into three distinct subtypes (Cd34-MC1, Cd34-MC2, and Cd34-MC3) as the mice aged (Figure 2c and d, Figure 2—figure supplement 3f). A comparative examination of their expression profiles highlighted the prevalence of pathways related to stem cell functions, including male gonad development, mesenchyme development, and regeneration, particularly within the Cd34-MC1 subtype (Figure 2e). Intriguingly, emerging markers such as Sox4 demonstrated distinctive expression in Cd34-MC1 at the adolescent stage (Figure 2f), indicating that adolescent Cd34-MC1 (Cd34^+^/Sox4^+^) may possess enhanced capabilities for differentiation, in contrast to Cd34-MC2 and Cd34-MC3. Immunofluorescence staining targeting Cd34 and Sox4 validated the observations regarding Cd34-MC1 (Figure 2g).

Additionally, by examining publicly available testicular scRNA-seq datasets from human infants and adults (GSE124263), we pinpointed a comparable CD34-MC subtype in both infantile and adult human testes (Figure 2h, Figure 2—figure supplement 3g-i). The pseudotime trajectories provided insights into the progenitor role of human CD34-MC population in LC lineage development (Figure 2i). Remarkably, these CD34-MC subtypes remain present within the human testicular interstitium through infancy to adulthood, manifesting unique transcriptional identities over time (Figure 2j and k).

### Significant glutathione reduction in interstitial cells during aging

To investigate transcriptional variations in the testicular interstitium with age, we conducted hierarchical clustering of cells (rows) based on the 2000 most variably expressed genes (columns) across different age categories. This approach identified two primary clusters of pseudotime-dependent genes with distinct signatures. The group of genes with increased expression in older samples included known aging markers like Spp1 and Cxcl10 (Figure 3a). Conversely, genes showing reduced expression in aging, such as Wnt5a and Kitl, formed a group potentially indicative of protective effects against aging (Figure 3a). Gene set enrichment analysis (GSEA) analysis revealed that genes with decreased expression in aging were significantly associated with metabolic pathways, including cholesterol biosynthesis and steroid hormones metabolism, suggesting age-related reduction in steroid hormone synthesis and metabolism (Figure 3b). Meanwhile, genes with increased expression were significantly linked to immune system pathways, like activation of innate and adaptive immunity, underscoring the prominence of immune mechanisms in the aging process of the testicular interstitium (Figure 3c).

**Figure 3.**
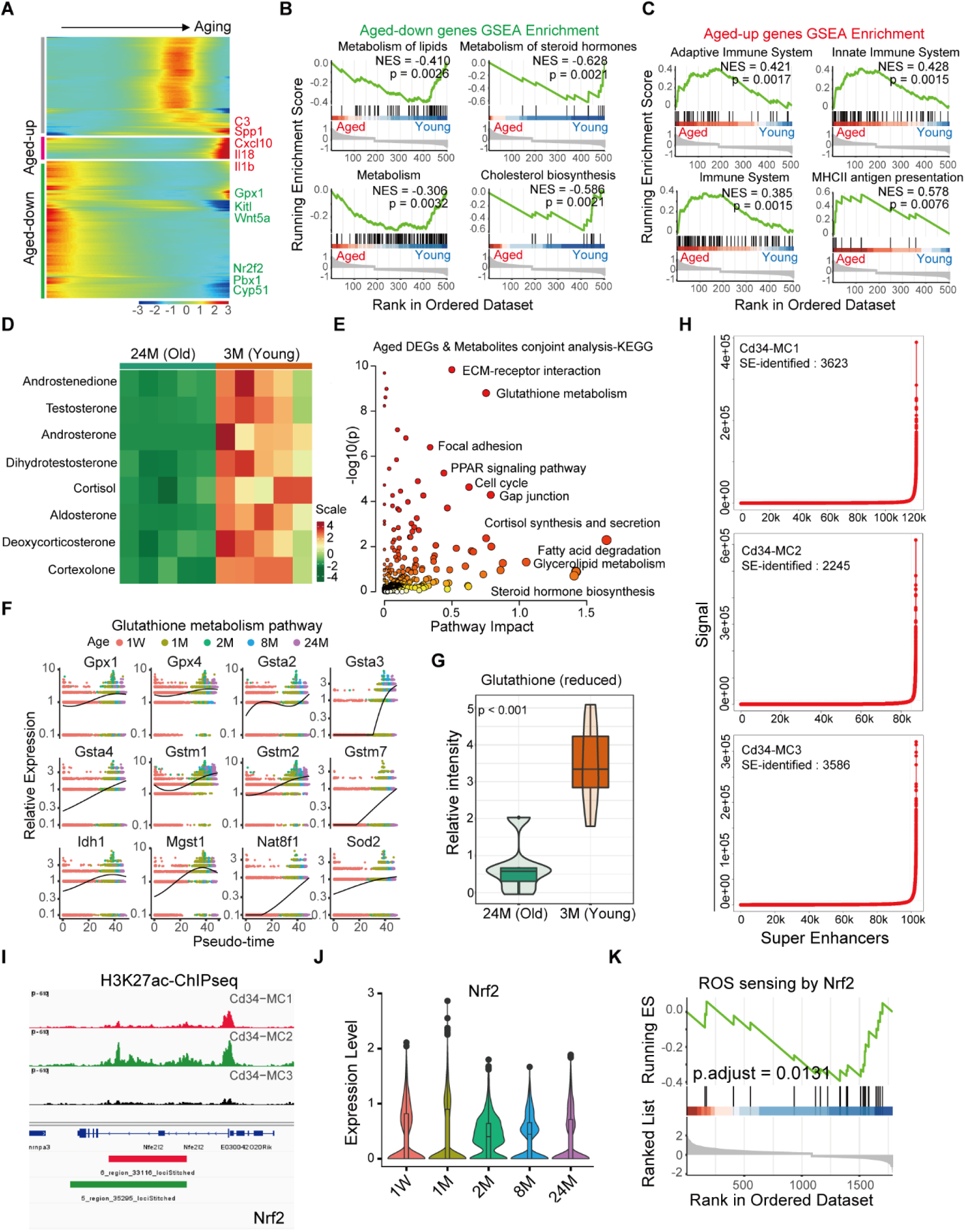
Alterations in glutathione metabolism within testicular interstitial cells during aging. (**a**) K-means clustering analysis identifies genes with differential expression across various populations of testicular interstitial cells with aging. (**b**) GSEA highlights metabolism-related pathways predominantly downregulated in older cells (p < 0.05). (**c**) GSEA reveals an enrichment of immune-related pathways in genes upregulated in the aged group (*p* < 0.05). (**d**) Targeted steroid hormone metabolomics analysis comparing the testicular lipid profile between young (3-month-old) and aged (24-month-old) mice (n=5). (**e**) Dot plot presents a combined pathway analysis, correlating upregulated metabolites with DEGs in older cells. (**f**) Trends in the expression of key genes within the glutathione metabolism pathway across the aging of interstitial cells. (**g**) Relative intensity of Glutathione (reduced form) in the testis derived from 3-month and 24-month-old mice (n=5, *p* < 0.01). (**h**) A ranked plot of Super-Enhancers (SEs) showcases those with the highest median H3K27ac scores across Cd34-MC1, Cd34-MC2, and Cd34-MC3 cells. (**i**) ChIP-seq profiles depict H3K27ac enrichment at Nrf2 gene loci. (**j**) Violin plot displays the expression trajectory of Nrf2 in Cd34-MC1 cells throughout aging. (**k**) GSEA indicates the influence of Nrf2 on the ROS sensing signaling pathway within Cd34-MC1 cells.

To elucidate the metabolic alterations associated with aging in interstitial cells, we analyzed testis samples from individuals aged 3 months and 24 months using untargeted metabolomics via Liquid Chromatography-Mass Spectrometry. Principal component analysis demonstrated general metabolic similarity across samples (Figure 3—figure supplement 4a), while a volcano plot highlighted significantly altered metabolites (Figure 3—figure supplement 4b). Notably, aged tissues exhibited increased levels of peroxides and hydroxylates, such as beta-Hydroxymyristic acid, D-a-Hydroxyglutaric acid, and 6-Hydroxycaproic acid (Figure 3—figure supplement 4c). Furthermore, spectrometry-based steroid metabolome profiling revealed a significant decrease in steroid hormones, particularly testosterone, in aged tissues (Figure 3d).

To examine the interplay between gene expression shifts and metabolic alterations, we merged data from single-cell transcriptomics with metabolomics. Utilizing MetaboAnalyst 5.0 for joint-pathway analysis, we identified that aging-associated DEGs and metabolites were significantly concentrated in pathways linked to glutathione metabolism (Figure 3e). We also noted an age-related progressive increase in the expression of glutathione oxidase encoding genes (Figure 3f, Figure 3—figure supplement 4d), while reduced form of glutathione decreased (Figure 3g). What’s more, cells in the old testis interstitium express most of the genes in the GenAge database, such as Gpx4, Gsta4 (Figure 3—figure supplement 4d, e)

To pinpoint crucial transcription factors (TFs) regulating the expression of genes involved in glutathione metabolism, we applied super-enhancer mapping using H3K27 acetylation (H3K27ac) Cut&Tag in Cd34-MCs (Figure 3h, Figure 3—figure supplement 4f). Notably, within Cd34-MC1 and MC2, super-enhancers associated with Nrf2 were identified. However, Nrf2 is no longer regulated by super-enhancers in Cd34-MC3 (Figure 3i). Protein-protein interaction (PPI) network analysis underscored that glutathione metabolism-related genes are potentially modulated by the oxidative stress-responsive transcription factor Nrf2 (Figure 3—figure supplement 4g). We observed that Nrf2 expression remains elevated from postnatal through adult stages, with a gradual decline in later life (Figure 3j). GSEA analysis implies that diminished Nrf2 expression in aged LCs may contribute to increased oxidative stress (Figure 3k).

### Unveiling Cd34-MC1 cells as potential LC progenitors through transplantation in LC-disrupted or failure mice

Subsequently, we evaluated the self-renewal capabilities and differentiation capacities of Cd34-MC cells sourced from mice at two distinct life stages: 2 weeks old (Cd34-MC1) and 24 months old (Cd34-MC2). Our findings revealed that Cd34-MC1 displayed enhanced self-renewal abilities and a higher clonogenic potential relative to their counterparts from older mice (Figure 4a). Furthermore, when cultured in a specialized medium, Cd34-MC1 cells from 2-week-old mice were capable of differentiating into LC lineages, whereas Cd34-MC2 cells, derived from 24-month-old mice, showed a markedly reduced capacity for proliferation and differentiation (Figure 4b).

**Figure 4.**
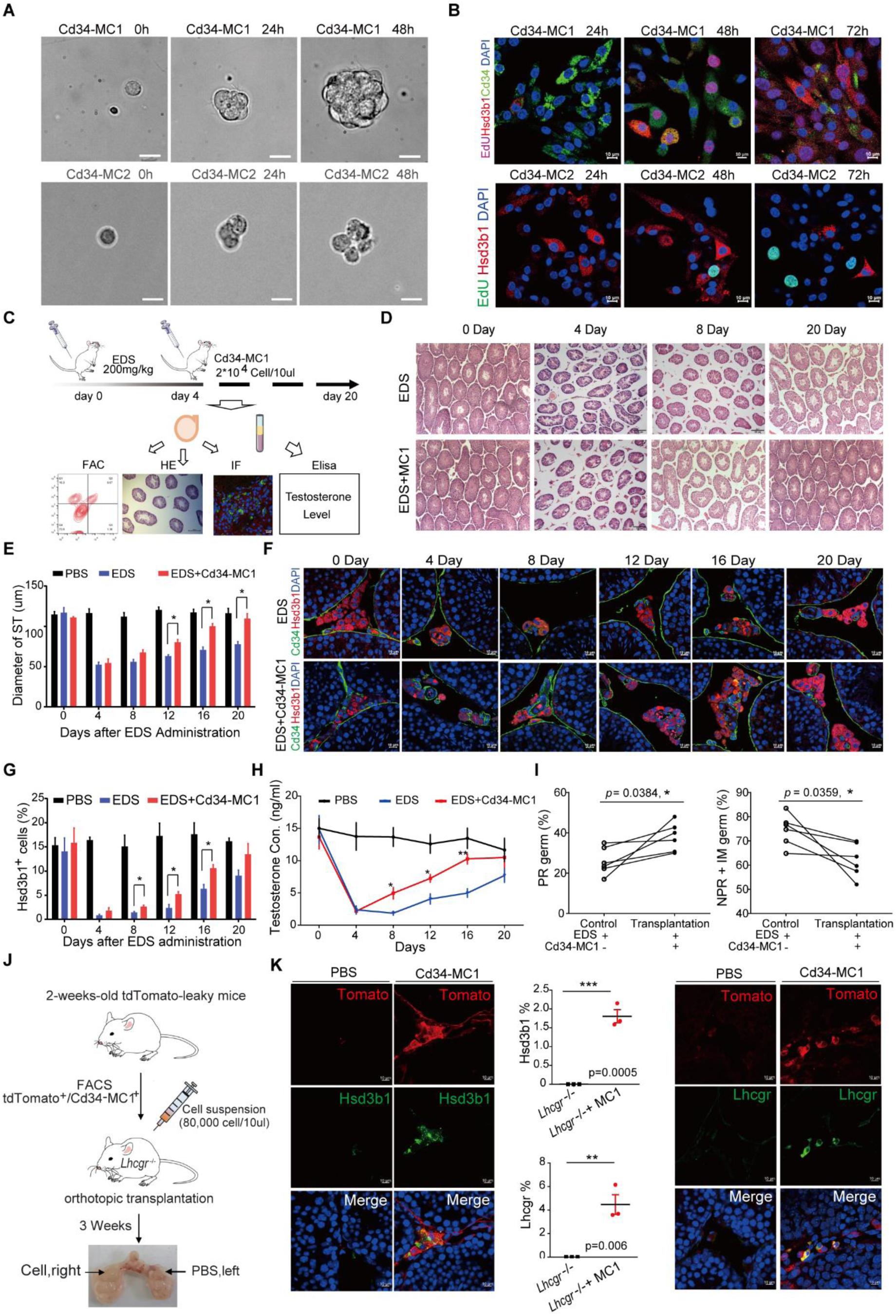
Enhancement of testosterone production in EDS-treated mice testes following Cd34-MC1 transplantation. (**a**) Bright-field microscopy images showing the capacity of clonal sphere formation between Cd34-MC1 and Cd34-MC2 cells originating from single cells. Scale bar: 10 µm. (**b**) Immunofluorescence staining showing the differentiation of Cd34-MC1 and Cd34-MC2 into LCs, marked by LC-specific marker expression. (**c**) Schematic diagram illustrating the experimental design for Cd34-MC1 cell transplantation. (**d**) Hematoxylin and eosin (H&E) staining of testis sections from control, EDS-treated, and and EDS-treated groups receiving Cd34-MC1 transplants. Scale bar: 100 µm. (**e**) Quantitative analysis of seminiferous tubule diameter in the two groups. Three sections per slide and three slides per mouse testis were counted. Data are presented as the mean ± SEM. *, *p* < 0.05; **, *p* < 0.01; ***, *p* < 0.001 (compared to the control group, n = 6). (**f**) Immunofluorescence staining showing the presence of Hsd3b1^+^ cells in the testicular interstitium post Cd34-MC1 transplantation. Scale bar: 50 µm. (**g**) Quantitative analysis of the number of Hsd3b1^+^ cells in the three groups. Data are presented as the mean ± SEM. ***, *p* < 0.001 (compared to the EDS-treated group, n = 6). (**h**) Serum testosterone levels comparison among the groups. Data are presented as the mean ± SEM. *, *p* < 0.05 (compared to the EDS-treated group, n = 6). (**i**) Sperm motility assessment in both transplanted and non-transplanted mice, with progressive (PR) and non-progressive (NPR) motility sperm counts shown as mean ± SEM. *, *p* < 0.05; **, *p* < 0.01 vs. EDS-treated, n = 6. (**j**) Schematic diagram outlining the experimental design for the transplantation of Cd34-MC1-tdTomato cells into *Lhcgr*^-/-^ mice. (**k**) Immunofluorescence staining revealing the presence of Lhcgr^+^ and Hsd3b1^+^ cells within the interstitial region of the *Lhcgr*^-/-^ mouse testes after Cd34-MC1 cell transplant.

To further assess the in vivo functional potential of Cd34-MC1 cells, we examined their capacity to differentiate into LCs and ameliorate testosterone production in a LC-disrupted model induced by ethylene dimethanesulfonate (EDS, 200 mg/kg, day 0). We isolated Cd34-MC1 cells from 2-week-old mice via fluorescence-activated cell sorting (FACS) (Figure 4—figure supplement 5a) and transplanted these cells into the testicular parenchyma of mice on day 4 post-EDS treatment. Serum and testicular samples were collected at intervals (0, 4, 8, 12, 16, 20 days post-EDS treatment) for subsequent analyses (Figure 4c). Remarkably, the transplanted Cd34-MC1 cells localized specifically to the testicular interstitium, where they rapidly contributed to tissue structural recovery (Figure 4d and e). Significantly, the proportion of Hsd3b1^+^ cells, indicative of LCs, notably increased in the testes receiving Cd34-MC1 cell transplants (Figure 4f and g).

To determine if mice receiving cell transplants mirrored this diurnal testosterone rhythm, we collected blood samples at six intervals over a 24-hour period, 20 days following transplantation. Our findings demonstrated that the transplanted group displayed a diurnal testosterone secretion pattern (Figure 4—figure supplement 5b). Importantly, Cd34-MC1 transplantation markedly enhanced testosterone levels in EDS-treated mice between days 8 and 16 (Figure 4h). Given the pivotal role of testosterone in driving meiosis and spermiogenesis to completion, we assessed the presence of the meiotic marker Sycp3 via immunofluorescence staining. A substantial increase in Sycp3-positive cells was noted in the transplanted group relative to the EDS-only group (Figure 4—figure supplement 5c and d). Furthermore, sperm quality, specifically progressive motility, showed significant improvement in the day 20 transplanted group compared to controls (Figure 4i, Figure 4—figure supplement 5e).

To deepen our understanding of CD34-MC1 cells’ contribution to LC regeneration, we utilized mouse lineage tracing models with inducible tdTomato expression and Lhcgr knockout mice, the latter presenting with a deficit in LC development. In some tdTomato mice, we noted generalized epifluorescence across the body, even in the absence of Cre recombinase activity, indicating leaky expression of the Tomato reporter (Figure 4—figure supplement 5f). We isolated Tomato^high^ Cd34-MC1 cells from 2-week-old Tomato-leaky testis Figure 4—figure supplement 5g). Subsequently, we injected Tomato^high^ Cd34-MC1 cells into the 1-month-old (adolescence) Lhcgr^-/-^ mice (Figure 4j). After three weeks, a significant presence of Tomato-labeled LCs, characterized by the expression of Hsd3b1 and Lhcgr, was identified within the testicular interstitium (Figure 4k, Figure 4—figure supplement 5h). This evidence underscores the pivotal role of Cd34-MC1 cells in directly fostering the development of the LC lineage in the testicular interstitium.

### Identifying Sox4 as the key transcription factor maintaining stemness in Cd34-MC1 cells

Through H3K27ac ChIP-seq, we pinpointed specific super-enhancers linked to essential transcription factor (TF) loci, including Sox4, within CD34 MC1 cells (Figure 5a). GO analysis indicates that the super-enhancers in Cd34-MC1 are enriched in terms related to stem cell development (Figure 5b). Through GSEA analysis, we highlighted Sox4’s critical function in preserving the stemness of Cd34-MC1 cells (Figure 5c). Notably, Sox4 expression levels were significantly elevated in the postnatal stage but decreased markedly as development progressed (Figure 5d). Similarly, the levels of H3K27ac at the Sox4 locus showed a gradual decline from Cd34-MC1 to Cd34-MC2/3 (Figure 5e).

**Figure 5.**
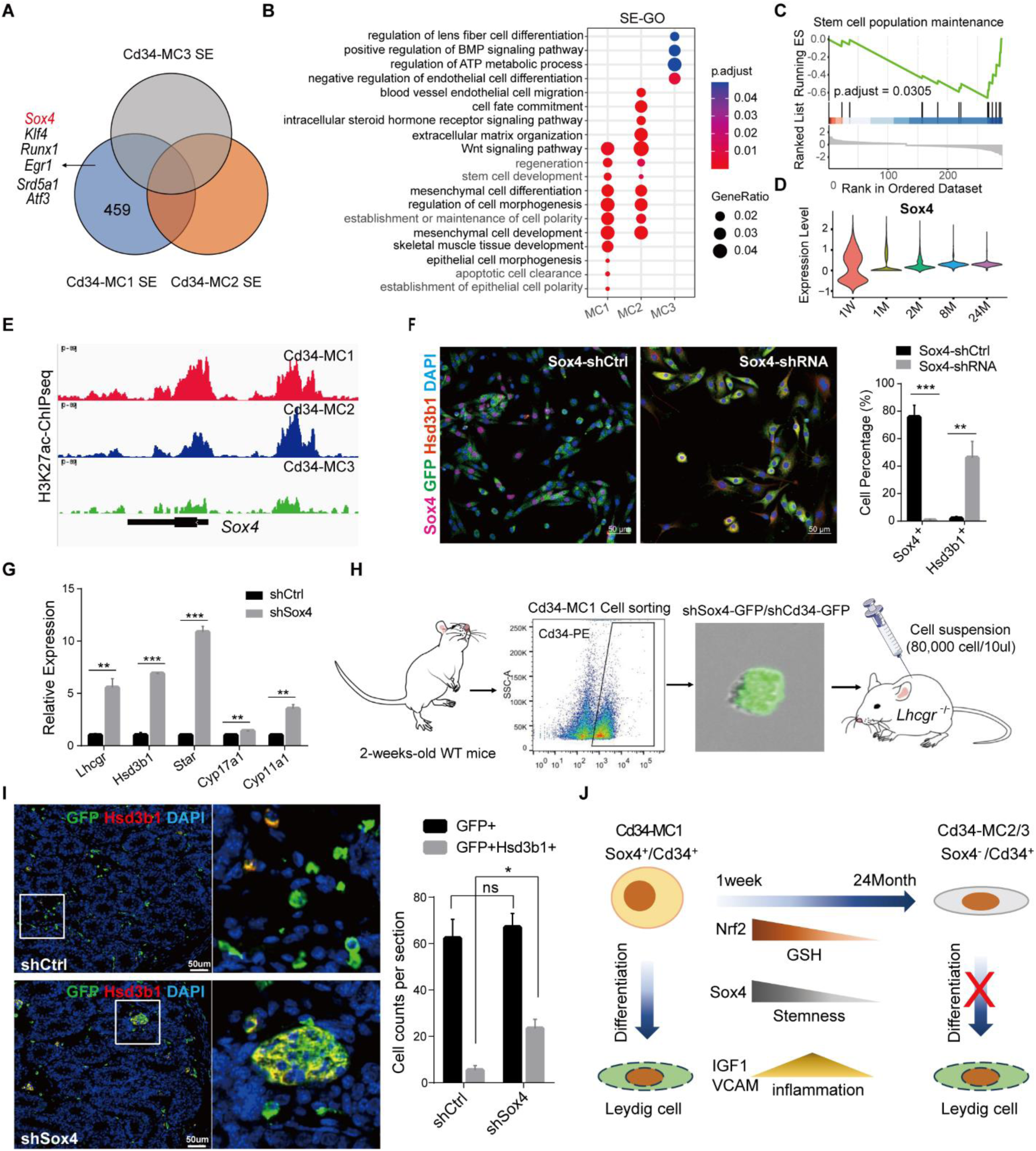
Super-enhancer (SE) landscape reveals Sox4 as a crucial transcription factor in maintaining stemness in Cd34-MC1. (**a**) Venn diagram displaying the specific SEs identified in Cd34-MC1. (**b**) GO analysis of the SEs in three subtypes of Cd34-MC. (**c**) GSEA showing the impact of Sox4 on pathways related to stem cell population maintenance. (**d**) Violin plots illustrating Sox4 expression across different developmental stages in Cd34-MC1. (**e**) ChIP-seq profiles highlight H3K27ac mark enrichment at the Sox4 locus within Cd34-MC1, Cd34-MC2, and Cd34-MC3. (**f**) Immunofluorescence staining identifies LCs expressing Sox4 shRNA (green), along with Sox4 (purple) and Hsd3b1 (red) positive cells in Cd34-MC1 after 48 hours of Sox4 knockdown. *, *p* < 0.05; **, *p* < 0.01; ***, *p* < 0.001. (**g**) qPCR analysis evaluates LC gene expression changes following 48 hours of Sox4 knockdown in Cd34-MC1 cells. *, *p* < 0.05; **, *p* < 0.01; ***, *p* < 0.001. (**h**) Schematic representation of Cd34-MC1 cell transplantation into *Lhcgr^-/-^* mice, with and without Sox4 knockdown. (**i**) Immunofluorescence staining shows Hsd3b^+^ LCs in the testicular interstitium of *Lhcgr^-/-^* mice post Cd34-MC1 cell transplantation. *, *p* < 0.05; **, *p* < 0.01; ***, *p* < 0.001. (**j**) Summary diagram illustrating the key findings of this research.

To explore the specific impact of Sox4 suppression, we utilized shRNA lentiviruses targeting Sox4 to achieve knockdown in primary Cd34-MC1 cells (Figure 5—figure supplement 6a and b). This manipulation led to a pronounced increase in Hsd3b1^+^ cell proportion after 48 hours of culture (Figure 5f). To gain deeper insights into Sox4 depletion’s effect on the transcriptional landscape, we conducted a quantitative PCR (qPCR) analysis on key mRNAs within the steroidogenic pathway. Sox4 knockdown notably upregulated expression of *Lhcgr*, *Hsd3b1*, *Star*, *Cyp17a1*, and *Cyp11a1* (Figure 5g). Additionally, the use of shRNA lentiviruses targeting Cd34 led to increased expression of *Lhcgr* and *Cyp17a1*, while decreasing the expression of *Hsd3b1*, *Star*, and *Cyp11a1* in Cd34-MC1 cells (Figure 5—figure supplement 6c-e). This indicates that *Cd34* is not necessary for the maintenance of cell stemness and cell differentiation.

Further, we injected Sox4 knockdown Cd34-MC1 cells or control cells into 1-month-old Lhcgr-/-mice (Figure 5h). Three weeks post-injection, the Sox4 knockdown group exhibited a substantial increase in LCs, marked by Hsd3b1 expression, within the testicular interstitium (Figure 5i). These results highlight the critical importance of Sox4 in preserving the stem cell lineage of Cd34^+^/Sox4^+^ MC1 cells.

## Discussion

The principal objective of this investigation was to elucidate the dynamics and diversity of LC across the lifespan, aiming to pioneer advancements in treatments for male infertility and promote reproductive health. Through comprehensive analyses, we have delineated distinct subpopulations of LC that derive from Cd34^+^/Sox4^+^ mesenchymal cells. Employing a combination of in vitro and in vivo approaches, our study reveals the remarkable regenerative properties of these specific cell types. Beyond cataloging LC heterogeneity, our work highlights the crucial role of metabolic rewiring and the impact of aberrant macrophage activity in the sustenance and function of LC (fig. 5J). Our results compellingly indicate that these elements are integral to LC homeostasis, with significant implications for male fertility.

Among our study’s most compelling discoveries was the delineation of Cd34^+^/Sox4^+^ mesenchymal cells as a pivotal progenitor source for LC maintenance and regeneration. Previous research has documented the presence of CD34^+^ mesenchymal cells within both fetal and adult human testes(Kuroda et al., 2004). Recent morphological studies have further uncovered CD34^+^PDGFRα^+^ telocytes in the interstitial spaces of mouse, rat, and human testes. These telocytes, forming intricate networks with LC, peritubular myoid cells, and vasculature(Abe, 2022), are known to express integrins α4, α9, β1, and VCAM1(elements suggestive of their role in establishing a stem cell niche conducive to LC differentiation). However, our findings spotlight Cd34^+^/Sox4^+^ mesenchymal cells as distinct stem/progenitor cells dedicated to LC maintenance and rejuvenation. Notably, these cells predominantly occupy peritubular regions in testes at one week postnatal, with their abundance significantly waning during later developmental phases. Moreover, our work revealed that Cd34^+^/Sox4^+^ mesenchymal cells from neonatal testes possess the capacity to form stem cell spheres and differentiate into LC. Impressively, when these cells are injected into adult testes experiencing LC disruption or failure, they foster regeneration within the interstitial compartment and elevate the count of Hsd3b1-positive cells. These insights affirm the critical function of Cd34^+^/Sox4^+^ mesenchymal cells in sustaining and restoring the testicular interstitium’s LC population, offering profound implications for understanding testicular physiology and developing treatments for male reproductive health issues.

While prior research has shed light on the morphological transformations of LC throughout postnatal development and aging(Guo et al., 2018; Guo et al., 2020; Li et al., 2017; Lukassen et al., 2018; Nie et al., 2022; Zhang et al., 2023), the complete spectrum of LC heterogeneity within the testicular interstitium over the lifespan remains partially unexplored. In our investigation, we performed an extensive analysis of LC across several postnatal phases, uncovering distinct clusters indicative of various LC subtypes. Among our findings, we identified a novel LC subpopulation AC, distinguished by the co-expression of Hsd3b1 and F4-80. Single-cell transcriptomic analysis unveiled distinctive gene expression patterns in AC, hinting at their potential role in immunological regulation within the testis. Remarkably, these cells were present in the testicular interstitium from childhood, possibly acting as a developmental link between Cd34^+^/Sox4^+^ mesenchymal cells and other LC clusters, as suggested by pseudotime analysis. Our research underscores the multifaceted roles of LC, extending beyond testosterone synthesis to include significant contributions to the immunological equilibrium within the testis.

Our study uncovers fascinating insights into the metabolic shifts within the testicular interstitium, building on prior work that identified aging LC as producing significantly more ROS in response to LH than their younger counterparts(Beattie et al., 2013). This increase in ROS suggests that an aging-related oxidative environment could play a role in the observed decline in testosterone production in older LC. We noted a reduction in intracellular antioxidants, like GSH, in LCs as they age. These antioxidants are essential for neutralizing cellular ROS and preventing oxidative damage to cellular functions. Indeed, earlier studies have linked GSH depletion with lowered testosterone production(Zirkin et al., 2008). Furthermore, our H3K27ac ChIP-seq analysis reveals that Nrf2—a pivotal transcription factor that activates antioxidant genes—is regulated by super-enhancers in Cd34-MC1 cells within the testicular interstitium. Intriguingly, Nrf2 expression in LC diminishes starting from adulthood. Prior research indicates that activating Nrf2 triggers a cascade of responsive genes that mitigate oxidative damage(He et al., 2020; Ma, 2013; Ngo and Duennwald, 2022). Therefore, we suggest that diminished Nrf2 expression in aging LC may lead to oxidative stress and a subsequent reduction in testosterone production. Further research is warranted to delve into how these metabolic alterations within cells affect LC functionality.

Our study also provides insights into the functional relationship between macrophages and LC subpopulations. Although the close association between testicular macrophages and LC is well-established, the specifics of their interaction have been less understood. Our data indicate that prepubertal macrophages emit factors like Vcam-1, Igf1, Visfatin, and Gas6, which are vital for LC development during the postnatal maturation of the testis. In contrast, as aging progresses, changes in macrophage polarization begin to adversely affect LC through several pathways, including cytokine environment alteration, provision of steroidogenic substrates, and modulation of oxidative stress. Our findings further suggest that aging testicular macrophages display significantly more immunosuppressive behavior compared to their counterparts in other tissues, a characteristic that can curtail LC steroidogenesis and potentially degrade overall testicular functionality. Moreover, these aged macrophages possess the ability to generate reactive oxygen species (ROS), potentially hindering Cd34-MC1 cell differentiation and impairing LC steroidogenesis. These insights highlight the critical role of macrophage regulation in maintaining LC function and underscore the detrimental effects of macrophage dysfunction on testicular health.

In summary, this study offers a significant contribution by mapping a detailed single-cell transcriptome atlas of cellular subpopulations within the murine testicular interstitium across postnatal development. Unveiling novel LC subpopulations and delineating their roles highlight the critical nature of cellular diversity in the testicular interstitium. Our insights into metabolic changes and the dynamic relationship between macrophages and LC pave the way for innovative treatments for age-associated testicular dysfunction and male infertility.

## Materials and Methods

### Animals

Wild-type C57BL/6j mice, Lhcgr knock-out mice in C57BL/6j background and Rosa26-CAG-loxp-stop-loxp-tdTomato mice were used in this study. The wild-type C57BL/6j mice were purchased from GemPharmatech, China, the Lhcgr knockout mice were donated by Zhenmin Lei’s laboratory, and the Rosa26-CAG-loxp-stop-loxp-tdTomato mice were donated by Zhao Meng’s laboratory.

### Isolation and capture of mouse testicular interstitial cells

Testis samples from C57BL/6j mice were dissected aseptically using sterilized scissors. The samples were washed three times with DPBS, and the tunica albuginea was carefully removed using sterilized tweezers to isolate the interstitial cells as our previous reported. The interstitial cells were then digested with 1 mg/ml collagenase type IV in DMEM-F12 (phenol red-free) at 37 °C for 10 minutes. During the digestion process, the tissues were gently pipetted up and down. The digestion process was halted by adding 2% FBS, and the resulting cell suspension was filtered through a 40-μm nylon mesh. The cells were then centrifuged at 350 rcf for 5 minutes at room temperature and suspended in DPBS or 2% FBS in DPBS for subsequent fluorescence-activated cell sorting (FACS) analysis.

### Flow cytometry analysis and cell sorting

For single-cell transcriptional profiling, the interstitial cells were suspended in DPBS containing 2% FBS and loaded onto a FACSAria III instrument for fluorescence-activated cell sorting. After sorting, the interstitial cells were selected for the construction of a single-cell RNA-sequencing library.

To identify markers and for cell culture experiments, testicular cells obtained after collagenase dissection were suspended in DPBS. The cells were incubated with primary antibodies, specifically Cd34-PE /Cd34-FITC and Cd36-APC, at 4 °C for 30 minutes to facilitate marker identification. To minimize fluorescence quenching, the samples were protected from light during this process. The unbound antibodies were removed by washing with DPBS containing 2% FBS. Subsequently, the cells were centrifuged at 350 rcf for 5 minutes at room temperature and re-suspended in DPBS containing 2% FBS for flow cytometry analysis and cell sorting.

### Clonal sphere formation assay for Cd34-MC1

Cd34^+^ cells were diluted to a density of 100 cells/ml and seeded into ultra-low adherent 96-well plates at a volume of 10 µl per well. A total of 150 µl of medium was added to each well. The medium consisted of DMEM/F12 supplemented with 5% chicken embryo extract, 1 nM dexamethasone, 1% ITS, 1% non-essential amino acids (NEAA), 1% N2 and 2% B27 supplements, 0.1 mM β-mercaptoethanol, and 1 ng/ml LIF, 20 ng/ml bFGF, 10 ng/ml EGF, 20 ng/ml PDGF-BB, and oncostatin-M.

Wells containing only one cell were marked, and the cultures were observed daily. Spheres were identified as free-floating spherical structures with a diameter greater than 50 µm. The cultures were maintained in a 5% CO2 water-jacketed incubator at 37 °C, and the medium was changed every 3 days.

### Cell differentiation culture

To initiate cell differentiation, 20,000 Cd34^+^ cells were suspended in a medium and plated on gelatin-coated glass-bottom cell culture dishes. The medium used was composed of phenol red-free DMEM/F12 supplemented with 2% FBS, 10 ng/ml PDGF-BB, 1 ng/ml LH, 1 nM TH, 70 ng/ml IGF1, and 1% ITS supplement.

The cells were incubated at 37 °C in a 5% CO2 environment for 3 days. Proliferation of the cells was assessed using the Click-iT Edu imaging kit, following the manufacturer’s instructions. Subsequently, differentiation was confirmed by immunostaining for lineage-specific markers of LCs.

### Cd34-MC1 cell transplantation

Ninety male C57BL/6j mice aged 3 months and ten male C57BL/6j mice aged 24 months were included in this study. The 3-month-old mice were randomly divided into three groups: Control group, EDS injury group, and Cell transplantation group (n=5 for each group at each time point). The control group received an intraperitoneal injection of DPBS, while the other two groups received a single dose of 200 mg/kg body weight of EDS, which led to the depletion of LCs in the adult testis within 4-6 days.

To assess whether the injection of Cd34^+^ cells could facilitate the recovery of LC dysfunction, the Cd34^+^ cells were washed with DPBS, approximately 20,000 Cd34^+^ cells in 10 µl of PBS were injected into the parenchyma of the recipient testes 4 days after the 3-month-old mice had received EDS treatment. Testes and serum samples from all animals were collected at 0, 4, 8, 12, 16, and 20 days after EDS treatment for subsequent flow cytometry analysis, histological analysis, and measurement of testosterone concentration.

### Cd34-MC1-tdTomato cell transplantation

Cd34-MC1-tdTomato cells were isolated from the testes of 2-week-old male tdTomato-leaky Rosa26-CAG-loxp-stop-loxp-tdTomato mice using flow cytometry. These isolated cells were then resuspended in DPBS. Subsequently, we performed orthotopic transplantations into the C57BL/6*^Lhcgr-/-^* mice. The right testis tissue of each C57BL/6*^Lhcgr-/-^* mouse received an injection of 80,000 Cd34-MC1-tdTomato cells in a volume of 10 μl per tissue, while the left testis tissue received a control injection of 10 μl of DPBS. After a three-week period, we collected testes from three of the mice that had undergone cell transplantation for immunofluorescence staining.

### Testosterone concentration assay

At each specified time point, cell culture supernatants and mouse sera were collected for the quantitative measurement of testosterone levels. The concentration of testosterone was determined using a commercially available ELISA kit, following the manufacturer’s instructions.

### Sox4-shRNA infection and assessment

We constructed shRNA sequences targeting Sox4, Cd34 and integrated them into the pTSBX-U6-shRNA-EF1 copGFP-2A-PURO lentiviral vector. Flow cytometry was utilized to sort Cd34-MC1 cells, which were then infected with lentiviruses carrying specific shRNA for 12 hours. After the infection period, cells were cultured in fresh medium for up to 48 hours. The efficiency of gene silencing was assessed using quantitative PCR (qPCR). The shRNAs and primers used in this paper is provided in **Supplementary Table S1**.

### Immunofluorescence of the mouse testis

The immunofluorescence stainings were performed on 5 μm formalin-fixed paraffin embedded (FFPE) sections from testis samples, rehydratation and heat-mediated antigen retrieval in 10 mM sodium citrate buffer solution (pH=6.0). After blocking with goat serum working solution for 30 minutes, individual sections were incubated overnight at 4 °C with a mix of diluted antibodies according to their recommended dilution ratio. Antigen detection was conducted using the appropriate combination of Alexa Fluor 488 (Thermo Fisher Scientific, A11034 and A11029) and 594 (Thermo Fisher Scientific, A11032 and A11037) secondary antibodies (1:500, respectively) for 2 hours at room temperature in the dark. All primary/secondary antibodies were diluted in 0.5%BSA. After three washes in PBS, sections were incubated with Hochest33342 (dilution 1:2000 in PBS) to facilitate nuclear visualization. Images were obtained under 20x or 63x oil objective with a Zeiss LSM780 Confocal Lase Scanning Microscope and analyzed using ZEN software. The primary antibodies and secondary antibodies used in this paper is provided in **Supplementary Table S2**.

### Histological examination

Testis were dissected from mice and fixative for 12 hours at 4 °C, embedded in paraffin, then sectioned into 5 μm slices. Before staining, tissue sections were dewaxed in xylene, rehydrated through decreasing concentration of ethanol, and washed in distilled water. Then the sections were stained with hematoxylin and eosin. After staining, sections were dehydrated through increasing concentration of ethanol and xylene, then mounted with neutral balsam. The sections were observed with Olympus BX63 microscope.

### Cut&Tag library construction

CUT&Tag for profiling histone H3K27ac was performed using Hyperactive Universal CUT&Tag Assay Kit for Illumina (vazyme, TD903-02), and anti-H3K27ac antibody were used to enrich target DNA fragments of Cd34-MC1, MC2 and MC3 cells obtained by flow sorting. TruePrep Index Kit for Illumina (v21.2, vazyme, TD202) were then used to DNA library construction.

### Data analysis of Cut&Tag

For H3K27ac-Cut&Tag data, clean reads were aligned to *Mus musculus* reference genome mm9 using bowtie2 software (v2.4.0). Duplicated reads were removed and only uniquely mapping reads were retained for further analysis. MACS2 (v2.1.1) was used to call peaks using the parameters: - p 0.05. For broad peaks, signal of H3K27ac from Cut&Tag data was used to define SEs. SEs were identified and annotated using the default parameters: -t 2500, according to ROSE algorithm(Whyte et al., 2013). The visualization of bigwig files was achieved using the IGV (v2.16.0) tool. Heatmaps and profiles of signal were plotted using deepTools.

### Quantitative real-time PCR (qRT-PCR)

Total RNA was extracted from the cells using the TRIzol reagent according to the manufacturer’s instructions. The concentration and purity of RNA were assessed using a NanoDrop spectrophotometer. Complementary DNA (cDNA) was synthesized from 1 μg of total RNA using the High-Capacity cDNA Reverse Transcription Kit. qRT-PCR was performed using the SYBR Green PCR Master Mix on the QuantStudio™ 5 Real-Time PCR System. The thermal cycling conditions were: 95°C for 10 minutes, followed by 40 cycles of 95°C for 15 seconds, and 60°C for 1 minute. Gene expression levels were normalized to GAPDH and calculated using the 2^-ΔΔCt^ method. Each sample was analyzed in triplicate.

### Processing of Single-cell RNA-seq data

Single-cell RNA sequencing data of testicular samples was aligned and quantified using the Cell Ranger Software Suite (v3.0.2, 10X Genomics). Firstly, we captured approximately 7,000 to 120,000 cells each sample. The reads were aligned to the *Mus musculus* reference genome mm10, and after alignment, filtering, barcode counting, and UMI counting using the cell count pipeline provided by Cell Ranger, feature-barcode matrices were generated. Secondly, we performed quality control using the Seurat R package(Hao et al., 2021; Qiu et al., 2017). Single cells that expressed less than 300 genes and more than 5,000 genes were excluded in case of the potential empty droplets and doublets, and genes expressed in fewer than 20 cells were excluded from further analysis. Individual cells that had less than 25% mitochondrial gene content were retained to remove the dead cells. Only eligible cells will be processed for downstream analysis. We use the Functions built into Seurat (v4.0)(Stuart et al., 2019). packages to correct for batching effects and merge data sets. The FindIntegrationAnchors function was used to find a set of anchors between five Seurat objects. These anchors can later be used to integrate the objects using the IntegrateData function. Thirdly, we performed nonlinear dimensional reduction using the Seurat package. The UMI count was normalized and then scaled based on 2,000 highly variable genes for the subsequent principal component analysis (PCA). We chose the top 10 PCs for the t-SNE and UMAP dimensional reduction. The FindClusters function with a resolution of 0.25 was used to identify a total of 17 distinct cell clusters, and using the FindMarkers command we identified the differentially expressed genes (DEGs) as well as the specific marker genes for each cell type.

### Gene enrichment analysis

To identify the functions and crucial pathways of differentially expressed genes across the different cell types, the DEGs were analyzed by the ClusterProfiler package (v4.0.5)(Wu et al., 2021; Yu et al., 2012) through the Gene Ontology (GO), Kyoto Encyclopedia of Genes and Genomes (KEGG) and Gene Set Enrichment Analysis (GSEA) databases. All the significantly enriched terms were determined using the p-value cutoff = 0.05, while the enrichment of GSEA terms also show the normalized enrichment score (NES), indicating statistical significance.

### Pseudotime analysis

To investigate the temporal progression and developmental trajectories of LC and Mesenchymal cell subtypes, we performed pseudotime analysis using Monocle (v2.20.0)(Trapnell et al., 2014). This analysis was conducted based on the expression patterns of highly variable genes that were identified by Seurat. Pseudotime represents a continuous trajectory that captures the inferred developmental progression of individual cells within a population.

### Definition of functional scores

In order to evaluate the functional characteristics of testicular macrophages at different stages of postnatal development, we employed the “AddModuleScore” function within Seurat to calculate functional scores for specific gene sets of interest. These gene sets were chosen based on their relevance to macrophage biology and included the following functional categories: macrophage migration (GO:1905517), antigen processing and presentation (GO:0019882), phagocytosis (GO:0006909), and macrophage cytokine production (GO:0010934). Additionally, we incorporated M1/M2 polarization gene sets, which were obtained from a previously published study(Sun et al., 2021).

### Cell-Cell interaction analysis

To investigate cell-cell interactions within the mouse testis, we utilized the CellChat software (v1.1.2)(Jin et al., 2021). Several functions within CellChat were employed to analyze and visualize the signaling pathways involved in cell-cell communication. First, we utilized the “netVisual_chord_gene” function to visualize the signaling pathways originating from macrophages and targeting LCs or Cd34^+^ mesenchymal cells. This allowed us to explore the interactions and potential signaling events between these cell types. Next, the “netVisual_aggregate” function was employed to display selected signaling networks related to the signaling pathways of interest. This function enabled us to focus on specific signaling pathways and investigate the interactions between different cell populations within the testis. Furthermore, the “netVisual_bubble” function was utilized to present specific ligand-receptor pairs associated with the signaling pathways of interest. Ligand-receptor pairs such as VCAM, IGF, VISFATIN, and GAS signaling pathways were specifically examined, providing insights into the potential molecular interactions and communication between cells involved in these pathways.

### Statistical Analysis

Quantitative results are presented as mean ±SEM. ANOVA was used for multiple comparisons; where significance was detected, posthoc testing was then carried out (Graphpad Prism). All other quantitative analyses were analyzed using unpaired Students t-tests. Statistical analyses are indicated in figure legends. Significant outliers were detected using Grubb’s test (Graphpad Prism). Significance was determined at *p* < 0.05.

## Author contributions

X. H. performed the experiments, analyze the scRNA-seq data, and wrote the manuscript. K. X. performed the animal experiments. M. Y. analyzed the scRNA-seq data and helped in revising the manuscript. M. J. interpreted the data and provided technical support. W. L. interpreted the data and revised the manuscript. Z. L. provided the Lhcgr KO mice and provide technical support. A.P.X. interpreted the data, revised the manuscript, and provided supervision. W. Z. conceived and designed the experiments, interpreted the data, wrote and revised the manuscript, and provided supervision.

## Data and materials availability

The sequencing data supporting the conclusions of this study have been deposited in the GEO database under the accession numbers GSE262415 for H3K27ac-Cut&Tag data and GSE262416 for Single-cell RNA-seq data.

## Ethical approval statement

All mice used were housed and cared for in accordance with ethical guidelines, and experimental protocols were approved by the ethics Committee of Sun Yat-sen University (No. 2017-133).

## Supplementary Materials

**figure supplement 1.**
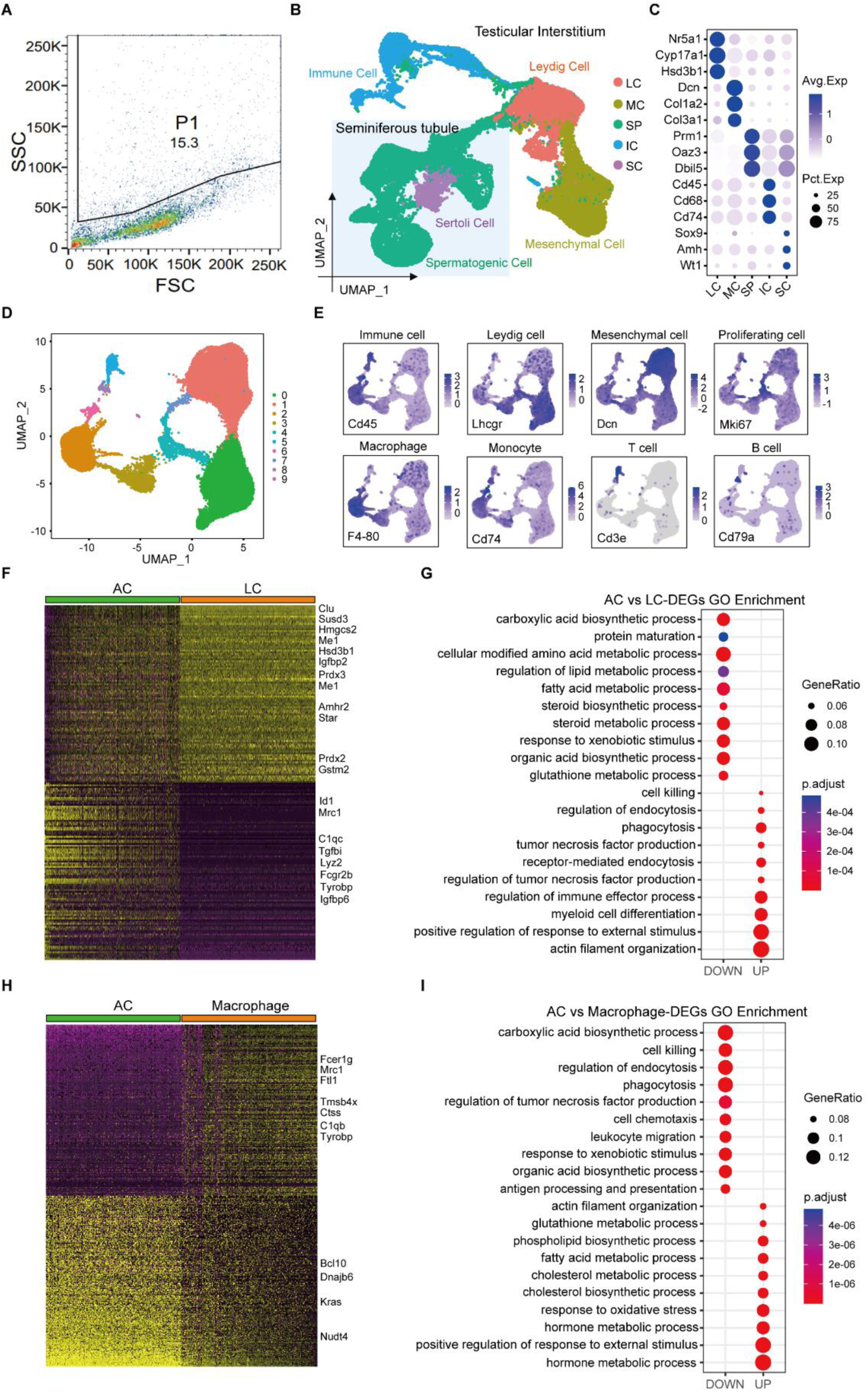
Single-cell transcriptome profiling of the mouse testicular interstitium. (**a**) Isolation of mouse testicular interstitial cells via fluorescence-activated cell sorting (FACS) for single-cell RNA sequencing (scRNA-seq). (**b**) A Uniform manifold approximation and projection (UMAP) visualization showing all identified cell types from scRNA-seq, with analysis focused exclusively on cells from the testicular interstitium. (**c**) Dot plot displaying characteristic DEGs across identified cell types. (**d**) UMAP representation highlighting distinct cell populations within the mouse testicular interstitium across neonatal, pre-pubertal, adolescent, adult, and aged stages in mice. (**e**) UMAP plot showing expression patterns of selected marker genes. (**f**) Heatmap presenting the differentially DEGs between AC and LC. (**g**) GO enrichment analysis of AC-upregulated and AC-downregulated DEGs (compared with LC). Gene expression levels are scaled and colored based on Z-scores, while enrichment significance is indicated by color and adjusted p-values. (**h**) Heatmap presenting the DEGs between AC and macrophages. (**i**) GO enrichment analysis of AC-upregulated and AC-downregulated DEGs (compared with macrophages). Gene expression levels are scaled and colored based on Z-scores, while enrichment significance is indicated by color and adjusted p-values.

**figure supplement 2.**
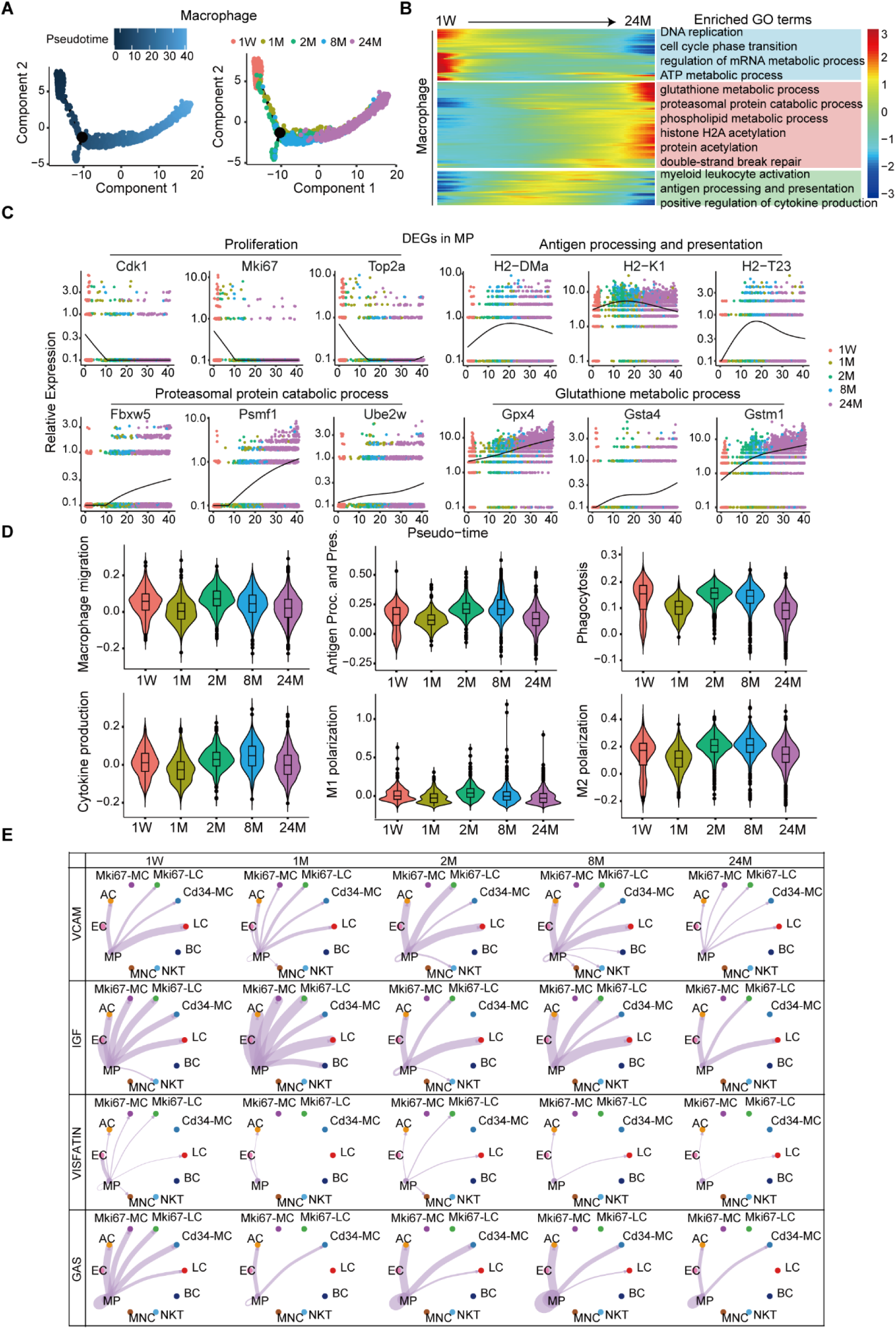
Dynamic functional changes of testicular macrophages across the postnatal lifespan. (**a**) Pseudotime trajectory analysis for macrophages across five developmental stages postnatally, depicted with distinct colors. (**b**) Heatmap illustrates temporal shifts in GO term enrichment for macrophages, mapped along the pseudotime continuum. (**c**) Expression trend plots for selected genes within macrophages across pseudotime, associated with processes like proliferation, antigen processing and presentation, proteasomal protein degradation, and glutathione metabolism. (**d**) Violin plots reveal changes in functional scores for macrophages during development, assessing aspects such as migration, antigen processing and presentation, phagocytosis, cytokine production, and M1/M2 polarization. (**e**) Circle diagrams depict key signaling pathways initiated by macrophages, highlighting VCAM, IGF, VISFATIN, and GAS pathways. Edge colors match the signaling sources, and the width of each edge denotes the likelihood of communication, with thicker lines indicating more robust signaling.

**figure supplement 3.**
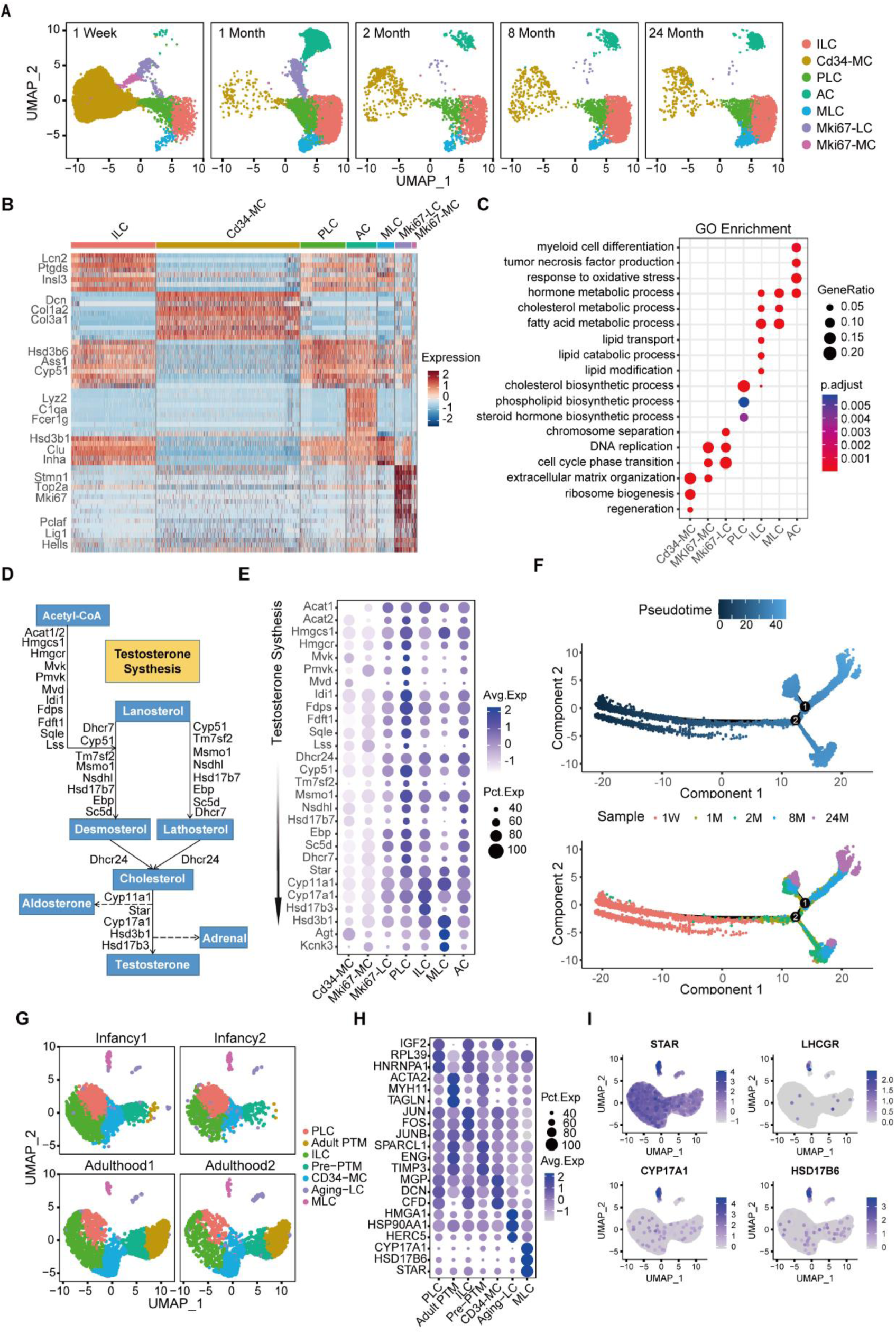
Subpopulation classification and identification within LC and MC across developmental stages. (**a**) UMAP visualization displays LC and MC subtypes across various developmental phases: neonatal (1 week), adolescent (1 month), adult (2 months), middle-aged (8 months), and aged (24 months). (**b**) Heatmap of the top ten differentially expressed genes for each LC and MC subtype, with expression levels normalized and depicted through a Z-score-based color gradient. (**c**) Gene Ontology (GO) enrichments for each LC subtype, with significance denoted by adjusted p-value color coding. (**d**) Schematic representation of testosterone biosynthesis, emphasizing genes implicated in the process. (**e**) Dot plot detailing gene expression relevant to testosterone synthesis across stages of LC lineage differentiation. (**f**) Pseudotime trajectory analysis delineates differentiation pathways from undifferentiated MCs towards mature LCs, showcasing expression evolution within the LC lineage from neonatal to aged stages. (**g**) UMAP analysis reveals cell type distribution within the testicular interstitium of infant (n=2) and adult (n=2) human testes, based on dataset GSE124263. (**h**) Dot plot illustrates marker gene expression profiles, identifying key cell types within the human testicular interstitium. (**i**) Expression analysis showing significant levels of STAR, LHCGR, CYP17A1, and HSD17B6 in mature human Leydig cells (MLC).

**figure supplement 4.**
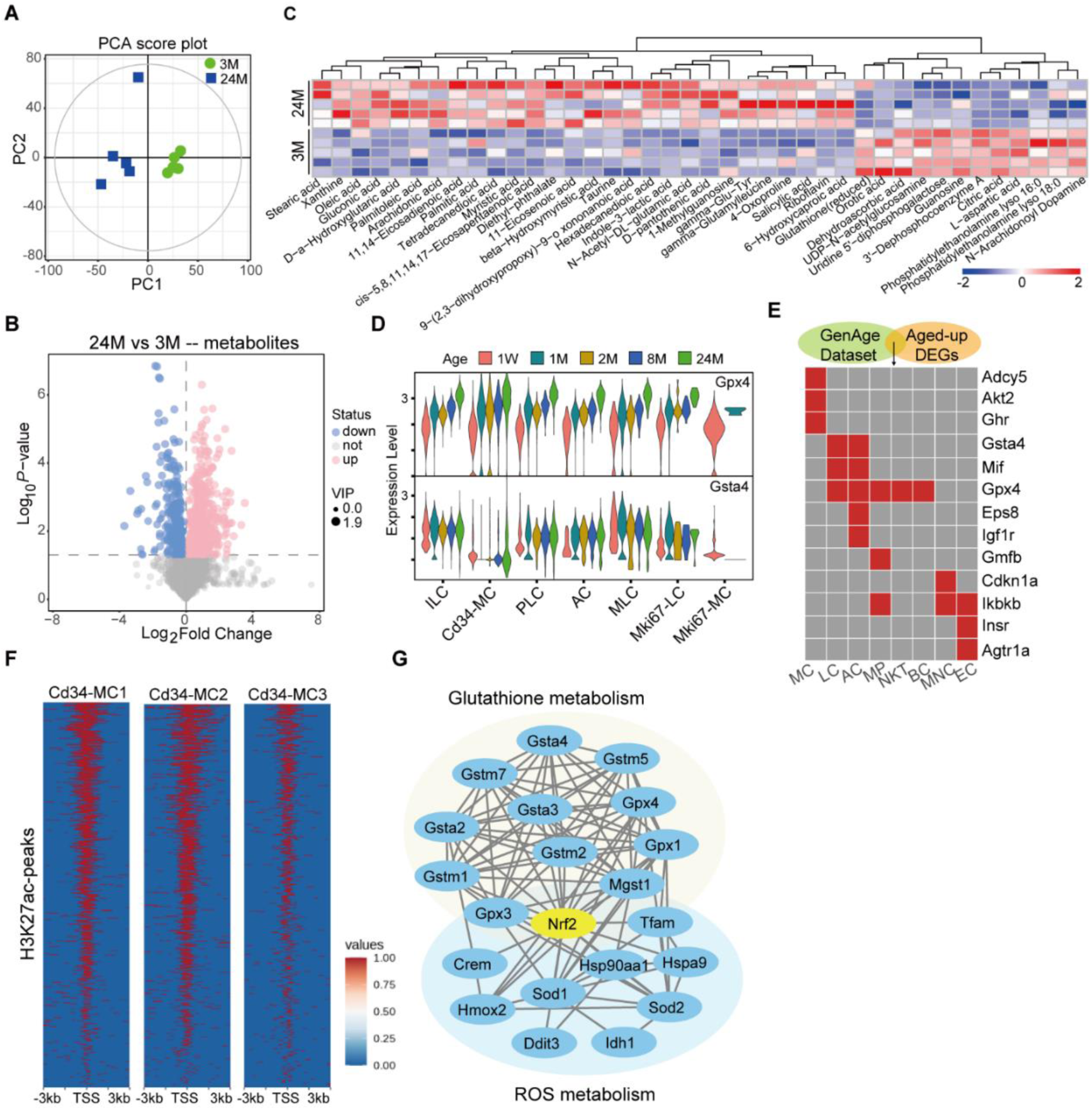
Transcriptional changes and metabolic alterations in the testicular interstitium during aging. (**a**) Principal Component Analysis (PCA) score plot demonstrating distinct metabolite profiles between the young and aged groups (n=5). (**b**) Volcano plot identifies differentially abundant metabolites comparing testes from 3-month-old and 24-month-old mice (n=5). (**c**) Heatmap showcases key metabolites that differ significantly between young and aged groups. (**d**) Violin plot illustrates *Gpx4* and *Gsta4* expression across specific subpopulations. (**e**) Heatmap shows that aging-related genes in the GenAge database are expressed in testicular subpopulations of aged mice. (**f**) The intensity of H3K27ac at the transcription start site (TSS) in Cd34-MC1, CD34-MC2, and Cd34-MC3 cells. (**g**) Protein-protein interaction (PPI) analysis emphasizes the involvement of Nrf2 in both glutathione and reactive oxygen species (ROS) metabolism pathways.

**figure supplement 5.**
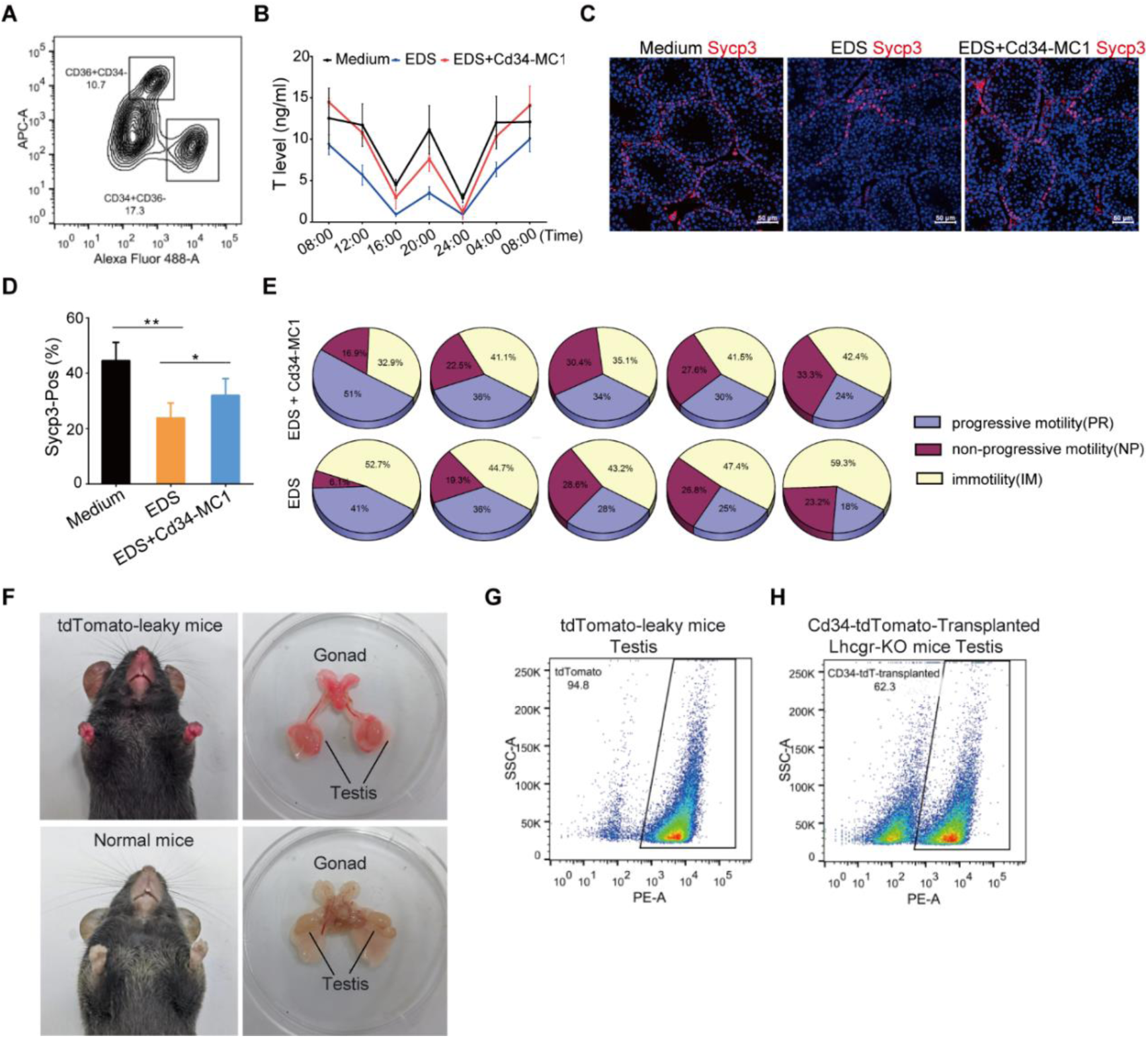
Restoration of testicular function by transplantation of Cd34-MC1 in LC-disrupted mouse models. (**a**) Flow cytometry results showing the percentage of Cd34^+^/Cd36^-^ (CD34-MC1) cells in the testes of 2-week-old mice. (**b**) Comparison of the diurnal testosterone secretion rhythm in control, ethylene dimethane sulfonate (EDS)-treated, and Cd34-MC1-transplanted EDS-treated groups. (**c**) Immunofluorescence imaging with anti-Sycp3 antibody to identify meiotic spermatocytes in the testes. (**d**) Quantitative analysis reveals a marked increase in Sycp3^+^ seminiferous tubules per section in Cd34-MC1-transplanted mice versus those receiving EDS treatment only. (**e**) Evaluation of sperm motility differences between Cd34-MC1-transplanted mice and their non-transplanted counterparts. (**f**) Comparative imagery of tdTomato-leaky mice against standard inducible tdTomato mice (uninduced). (**g**) FACS-based isolation of Cd34-MC1 cells from 2-week-old tdTomato-leaky mice for subsequent transplantation. (**h**) Flow cytometry assessment revealing the presence of tdTomato^+^ cells in the testes of Lhcgr knockout mice.

**figure supplement 6.**
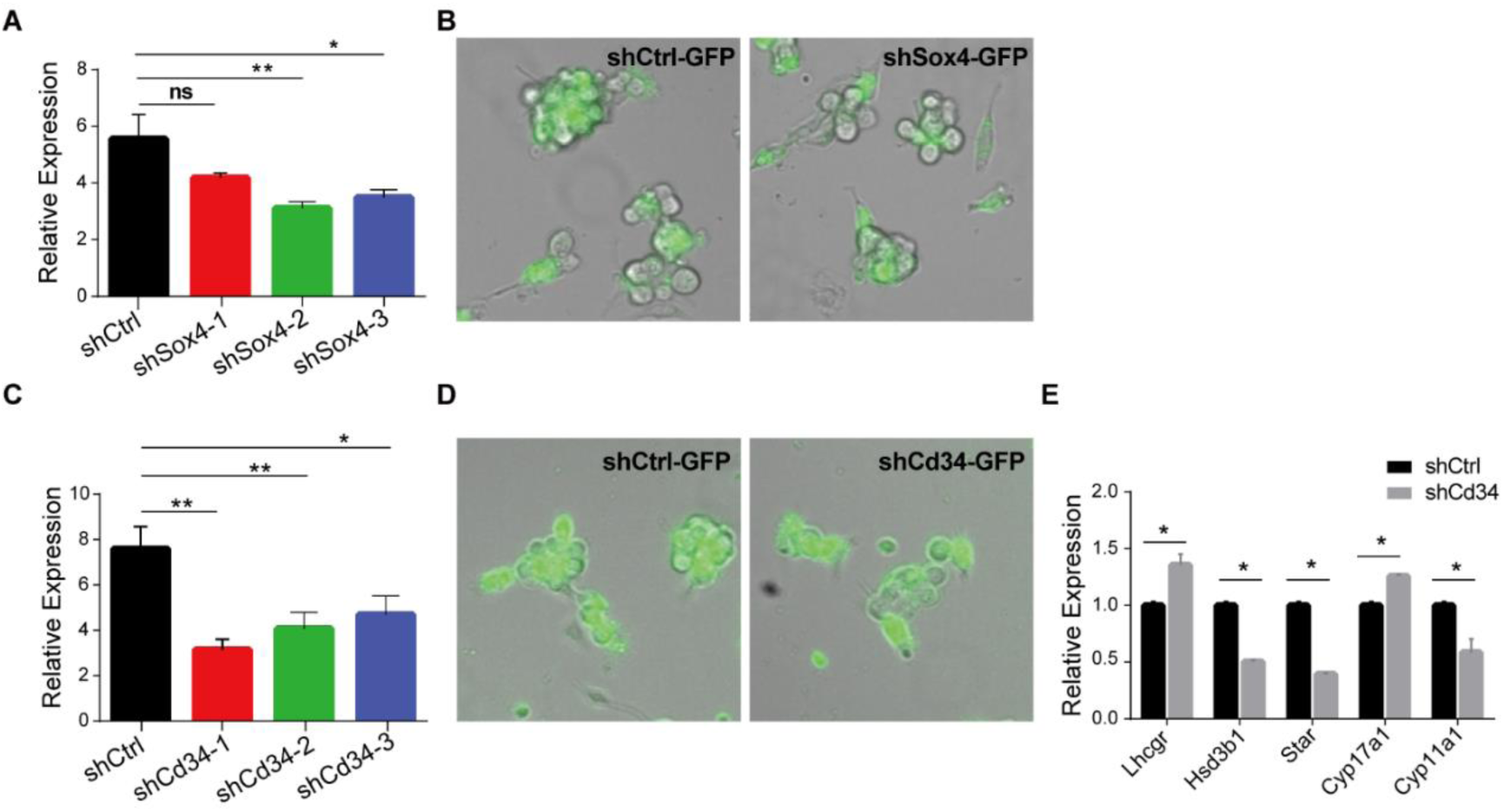
Master transcription factor Sox4 governing the gene expression program of Cd34-MC1. (**a**) qPCR analysis demonstrating the knockdown efficiency of Sox4 shRNAs in Cd34-MC1 cells. (**b**) Fluorescence indicating the infection efficiency of Sox4 shRNA lentiviruses in Cd34-MC1 cells. (**c**) qPCR analysis demonstrating the knockdown efficiency of Cd34 shRNAs in Cd34-MC1 cells. (**d**) Fluorescence indicating the infection efficiency of Cd34 shRNA lentiviruses in Cd34-MC1 cells. (**e**) qPCR analysis revealing alterations in Leydig cell gene expression 48 hours after Cd34 knockdown in Cd34-MC1 cells.

### Supplementary Tables

**Table S1.**
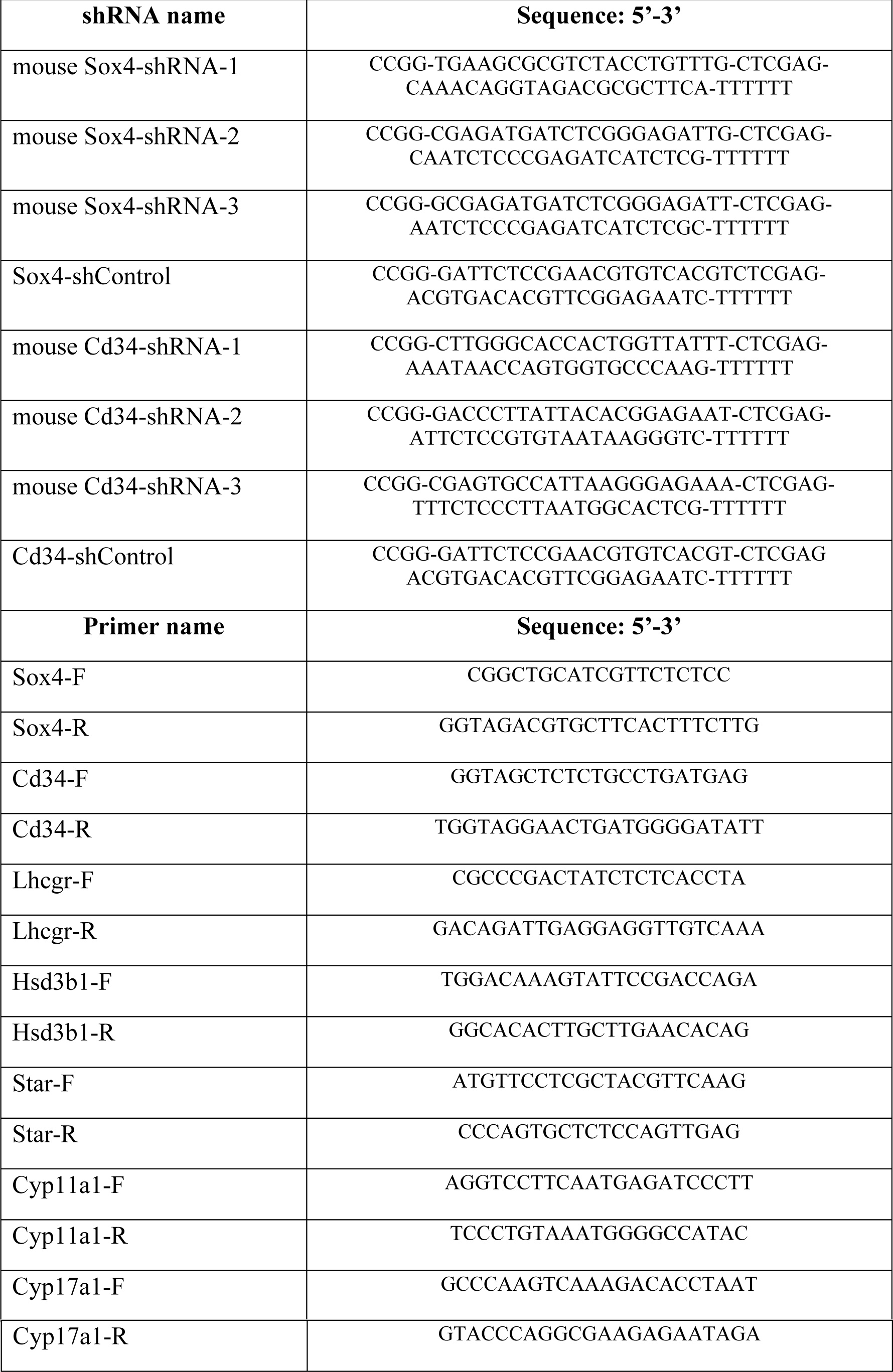
shRNAs and primers.

**Table S2.** Primary and secondary antibodies.

| **Primary antibodies** |  |  |
| --- | --- | --- |
| anti-CD34 | Abcam | Cat# ab81289 |
| anti-Adgre1(F4-80)-Rabbit | Cell Signaling Technology | Cat# 30325T |
| 3β-HSD (A-1) | Santa Cruz Biotechnology | Cat# sc-515120 |
| anti-Sycp3 | Santa Cruz Biotechnology | Cat# sc-74569 |
| anti-Lhcgr | Santa Cruz Biotechnology | Cat# sc-393592 |
| anti-Sox4 | Abcam | Cat# ab-243739 |
| anti-mouse CD34-FITC | BD | Cat# 553733 |
| anti-mouse CD36-APC | BioLegend | Cat# 102612 |
| anti-mouse CD34-PE | BioLegend | Cat# 128610 |
| Anti-Histone H3 (acetyl K27) | abcam | Cat# ab4729 |
| **Secondary antibodies** |  |  |
| Goat Anti-Rabbit AF594 | Thermo Fisher Scientific | Cat# A11037 |
| Goat Anti-Rabbit AF488 | Thermo Fisher Scientific | Cat# A11034 |
| Goat Anti-Mouse AF594 | Thermo Fisher Scientific | Cat# A11032 |
| Goat Anti-Mouse AF488 | Thermo Fisher Scientific | Cat# A11029 |

**Table S3.** Chemicals, Peptides, and Recombinant Proteins.

|  |  |  |
| --- | --- | --- |
| DPBS | Thermo Fisher Scientific | Cat# 14190-144 |
| Fetal Bovine Serum | Thermo Fisher Scientific | Cat# 9002-93-1 |
| Type IV Collagenase | Thermo Fisher Scientific | Cat# 17104019 |
| Hoechst 33342 | Thermo Fisher Scientific | Cat# H21492 |
| DMEM/F-12 (phenol red-free) | Thermo Fisher Scientific | Cat# 21041025 |
| Penicillin-Streptomycin | Thermo Fisher Scientific | Cat# 15140-122 |
| KnockOut Serum Replacement | Thermo Fisher Scientific | Cat# 10828-028 |
| NEAA | Thermo Fisher Scientific | Cat# 11140050 |
| Insulin-Transferrin-Selenium | Thermo Fisher Scientific | Cat# 41400045 |
| GlutaMAX | Thermo Fisher Scientific | Cat# 35050061 |
| N2 | Thermo Fisher Scientific | Cat# A1370701 |
| B27 | Thermo Fisher Scientific | Cat# A1486701 |
| β-mercaptoethanol | Merck | Cat# ES-007-E |
| BSA | AMRESCO | Cat# 0332-100G |
| Gelatin | AMRESCO | Cat# 9764 |
| Triton X-100 | AMRESCO | Cat# 9002-93-1 |
| 0.25%Trypsin | Corning | Cat# 25-053-CI |
| Recombinant mouse PDGF-BB | PeproTech | Cat# 315-18-10 |
| Recombinant mouse LIF | PeproTech | Cat# 250-02-25 |
| Recombinant mouse bFGF | PeproTech | Cat# 450-33-50 |
| Recombinant mouse TGF-β | PeproTech | Cat# 100-21-2 |
| Recombinant mouse IGF1 | PeproTech | Cat# 100-11-100 |
| Recombinant murine EGF | PeproTech | Cat# 315-09 |
| Luteinizing hormone (LH) | PeproTech | Cat# 12-4208 |
| Oncostatin M | R&D | Cat# 28-11178-2 |
| 3,3',5'-Triiodo-L-thyronine (TH) | Sigma | Cat# T2877 |
| Dexamethasone | Sigma | Cat# D4902 |
| 4% paraformaldehyde, PFA | Dingguo, China | Cat# AR-0211 |
| Ethylene dimethanesulfonate | Shaoyuan, China | Cat# EN300-137131 |
| Chicken Embryo Extract | absin | Cat# abs80002 |
| Gentian Violet solution | Sigma-Aldrich | Cat# 49144 |
| Goat Serum for Blocking | Beijing ZhanShanJinQiao | Cat# ZLI-9022 |
| Hematoxylin | Beijing ZhanShanJinQiao | Cat# ZLI-9610 |
| Eosin | Beijing ZhanShanJinQiao | Cat# ZLI-9612 |

**Table S4.** Software and Algorithms.

|  |  |  |
| --- | --- | --- |
| FlowJo (v10.0.7) | Flowjo, LLC | https://www.flowjo.com/ |
| GraphPad Prism 6 | GraphPad Software Inc. | https://www.graphpad.com/scientific/software/prism/ |
| ZEN | Carl Zeiss | https://www.zeiss.com/ |
| SCA® CASA System (for Sperm analysis) | MICROPTIC | https://www.micropticsl.com/products/sperm-class-analyzer-casa-system/ |
| Cell Ranger (v3.0.2) | 10x Genomics | https://support.10xgenomics.com/singlecell-gene-expression/software/pipelines/latest/what-is-cell-ranger |
| R software (v4.0.3) | CRAN (Open Source) | https://cran.r-project.org/ |
| Seurat (v4.0) | Satija Lab | https://satijalab.org/seurat/ |
| Monocle (v2.20.0) | Trapnell et al., 2014 | http://cole-trapnell-lab.github.io/monocle-release/ |
| Cellchat | Suoqin Jin et al., 2021 | https://github.com/sqjin/CellChat |
| ClusterProfiler (v4.0.5) | Yu G, et al., 2012 | https://bioconductor.org/packages/release/bioc/html/clusterProfiler.html |
| ggplot2 (v3.3.5) | Wickham, et al., 2009 | https://ggplot2.tidyverse.org/ |
| ROSE | Richard A. Young, et al., | https://github.com/stjude/ROSE |
|  | 2013 |  |
| MetaboAnalyst 5.0 | Zhou G, et al., 2021 | https://www.metaboanalyst.ca/MetaboAnalyst/ModuleView.xhtml |

**Table S5.** Critical Commercial Assays.

|  |  |  |
| --- | --- | --- |
| CellTrace™ CFSE Cell Proliferation Kit | Thermo Fisher Scientific | Cat# C34554 |
| CLICK-IT EDU IMAGING KIT | Thermo Fisher Scientific | Cat# C10086 |
| Testosterone ELISA Kit | Beyotime, China | Cat# PT872 |

**Table S6.**
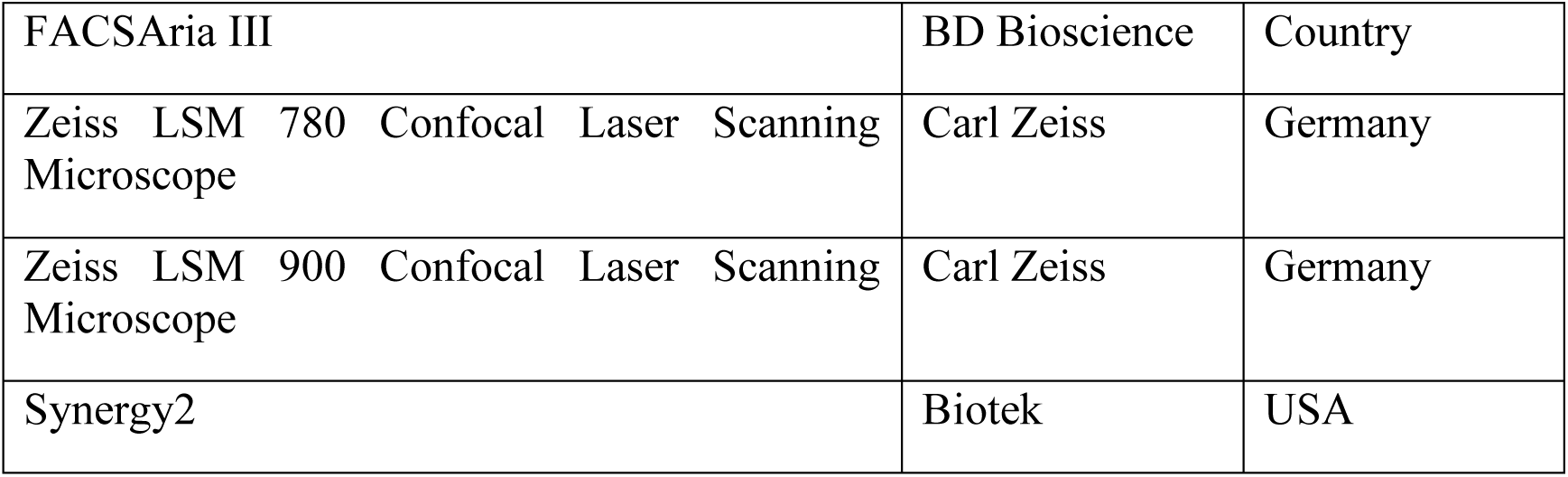
Instrument and Equipments.

|  |  |  |
| --- | --- | --- |
| FACSAria III | BD Bioscience | Country |
| Zeiss LSM 780 Confocal Laser Scanning Microscope | Carl Zeiss | Germany |
| Zeiss LSM 900 Confocal Laser Scanning Microscope | Carl Zeiss | Germany |
| Synergy2 | Biotek | USA |

